# Teneurin-2 at the Synapse Construction Site is a Signpost for Cargo Unloading from Motor Proteins

**DOI:** 10.1101/2022.09.13.507723

**Authors:** Sotaro Ichinose, Yoshihiro Susuki, Ryosuke Kaneko, Hirohide Iwasaki

## Abstract

In mature neurons, excitatory synapses are formed on the dendritic spine, whereas inhibitory synapses are formed on the dendritic shaft. Thus, it is primarily the accumulation of synaptic proteins that characterizes inhibitory synapses as distinct from non-synaptic regions. Protein accumulation is achieved by a combination of microtubule (MT)-based transport by kinesins and lateral diffusion across the plasma membrane; however, how and when proteins are released from kinesins remains unclear. Using primary cultured hippocampal neurons, we found that Teneurin-2 (TEN2) promotes synaptic protein accumulation by recruiting MTs via the representative MT plus end-tracking protein, EB1. MTs recruitment was enhanced when the extracellular domain of TEN2 successfully chose partners, and the lateral diffusion of TEN2 was inhibited. Conversely, if TEN2 partner choice is not achieved, MTs are not recruited, and thus synaptogenesis is not followed. Our study revealed that cargo release from kinesins through TEN2-MTs interactions supports the continuity from partner choice to synaptogenesis, which is a critical step in synaptic maturation.

## Introduction

Neurons connect to each other via synapses to generate dense neural circuits, and exchange information via synaptic receptors to properly perform brain activities that underlie various physiological functions. To this end, axons find their way to appropriate target regions (axon guidance), select appropriate synaptic partners within those regions (synaptic specificity), and form functional synapses (synaptogenesis) (Sanes & Zipursky, 2020). There are two types of synapses, excitatory and inhibitory, and their coordination, known as the E/I balance, allows for accurate information transfer; disruption of this balance can lead to disorders such as autism spectrum disorder and schizophrenia. These two types of synapses differ significantly in their cytoskeletons. Excitatory postsynapses form and mature on characteristic structures called spines, which are composed predominantly of actin, and microtubules (MTs) enter almost only in an activity-dependent manner (Gu et al., 2008; Hu et al., 2008; Jaworski et al., 2009; McVicker et al., 2016). In contrast, inhibitory synapses form directly on the dendritic shaft with both actin and MTs in the vicinity. Thus, at present, only the accumulation of inhibitory synapse-specific proteins distinguishes inhibitory postsynaptic membranes from non-synaptic membranes. At inhibitory synapses in the hippocampus, the primary receptors are GABA_A_ and glycine receptors, which are transported by KIF5 and KIF21, together with the scaffolding protein gephyrin (Labonté et al., 2014; Nakajima et al., 2012; Twelvetrees et al., 2010). After transport along MTs, receptors are exocytosed and move to the postsynaptic region via lateral diffusion (Dahan et al., 2003; Gu et al., 2016). At the postsynapse, receptors bind to actin filaments via gephyrin (Charrier et al., 2006; Giesemann et al., 2003). This prevents the diffusion of receptors and retains them in the post-synaptic region. Unnecessary receptors are endocytosed and transported by dynein (Fuhrmann et al., 2002; Kittler et al., 2000). Thus, the accumulation of receptors during synaptogenesis is a universal feature that identifies inhibitory postsynapses, and is supported by the spatiotemporal diversity of cytoskeleton-associated molecules.

In addition, the diversity of adhesion molecules is an important issue in synapses. Many studies have shown that numerous adhesion molecules belonging to diverse families are important for synaptic specificity (Sanes & Zipursky, 2020). However, it is not well understood whether synaptogenesis is continuously regulated by molecules that have achieved synapse specificity, or whether another regulatory mechanism is at work. Teneurin-2 (TEN2) is one of the few molecules that has been suggested to function in a continuity from synaptic specificity to synaptogenesis by the same molecule (Li et al., 2018). Two alternative splicing forms (SS+/SS-) exist in this molecule, with and without a 7 amino acid insertion: SS+ is involved in the formation of inhibitory synapses and SS- in the formation of excitatory synapses. At nascent excitatory synapses, presynaptic SS exhibits specificity by binding to postsynaptic Latrophilin-2/3 (Sando et al., 2019). Latrophilin is a G protein-coupled receptor that provides continuity from synaptic specificity to synaptogenesis by maturing synapses via downstream cAMP signaling (Sando & Südhof, 2021). In contrast, SS+ has been shown to potentially bind to an unknown binding partner other than latrophilin at nascent inhibitory synapses, but the process by which it matures inhibitory synapses is not well understood. An important consideration is that synaptogenesis does not occur spontaneously. The continuity from synaptic specificity to synaptogenesis arises only after immobilization via interaction with the binding partner. Conversely, when specificity is not achieved and the adhesion molecule diffuses laterally, synaptogenesis does not occur. Based on this idea and the spatiotemporal diversity of cytoskeleton-associated molecules at inhibitory synapses, we tested the following hypothesis. Upon successful partner selection, TEN2 was restricted to lateral diffusion by interacting with its binding partners. In parallel, MTs are captured near inhibitory synapses by TEN2, facilitating the release of receptors from motor proteins. This mechanism should contribute significantly to receptor accumulation during early development when the distribution of signaling molecules is inadequate.

In this study, we showed that TEN2 promoted receptor accumulation by recruiting MTs. TEN2 tends to localize to MTs-rich inhibitory postsynapses and is located at the semi-periphery approximately 85 nm away from the center of the postsynapse. Live imaging showed that MT dynamics were significantly reduced when TEN2 lateral diffusion was inhibited, suggesting that TEN2 binds to its binding partners and traps MTs. Accumulation of gephyrin and GABA_A_ receptor γ2 subunits was reduced in TEN2 knockdown neurons, suggesting that TEN2 is essential for the maturation of inhibitory postsynaptic synapses. Furthermore, overexpression of the MT-binding domain of TEN2 suppressed gephyrin clustering via a dominant-negative effect. These results suggest continuity from synaptic specificity to synaptogenesis, in which receptor accumulation is promoted only when TEN2 successfully selects a partner. This continuity is supported by a molecular basis in which TEN2 functions as a relay point for intracellular transport by providing an “end point” for kinesin transport and an “entry point” to the synapse for lateral diffusion.

## Results

### Cluster analysis of inhibitory postsynapses and correlation with TEN2

Excitatory postsynapses form and mature on characteristic structures called spines, which are composed predominantly of actin, and MTs enter almost only in an activity-dependent manner. In contrast, inhibitory synapses form directly on the dendritic shaft with both actin and MTs in the immediate vicinity. Thus, at present, only the accumulation of inhibitory synapse-specific proteins distinguishes inhibitory postsynaptic membranes from non-synaptic membranes. The accumulation of receptors is supported by the spatiotemporal diversity of cytoskeleton-associated molecules (Charrier et al., 2006; Fuhrmann et al., 2002; Giesemann et al., 2003; Gu et al., 2016; Kittler et al., 2000). However, the extent to which diversity exists is not yet fully understood. Therefore, we decided to observe and classify the cytoskeletal states of the inhibitory postsynapses. In this study, we defined postsynapses as those with immunofluorescence staining of gephyrin intensity above a certain value (Figure 1–figure supplement 1A-D).

To visualize the diversity of the cytoskeleton at inhibitory post-synapses, we quantified the intensity of MTs and actin at 20 days in vitro (DIV 20; Figures 1A and B). Dendritic MTs were visualized using MAP2 antibodies. After processing of MAP2 and actin intensity in the postsynaptic area in two dimensions using the uniform manifold approximate projection (UMAP) method, it was found appropriate to divide the data into three clusters by hierarchical clustering (Figure 1–figure supplement 1E and F). Cluster 1 had high MAP2 intensity, and cluster 3 had high actin intensity. Cluster 2 was considered to have low levels of both MAP2 and actin (Figure 1C). Applying findings from previous literature regarding the function of cytoskeleton-related molecules, we next inferred that the clustering results reflect the following cellular functions: Cluster 3 belongs to stable postsynapses anchored by gephyrin and actin (Charrier et al., 2006; Giesemann et al., 2003); cluster 1 belongs to dynamic postsynapses where receptors are being brought in and out by an MTs-based transport system (Labonté et al., 2014; Nakajima et al., 2012; Twelvetrees et al., 2010); cluster 2 belongs to intermediates where intense lateral diffusion is occurring (Dahan et al., 2003; Gu et al., 2016). Transitions between clusters are thought to occur in a developmental stage- and activity-dependent manner, similar to that in excitatory synapses. However, the transition between clusters 1 and 2; that is, how receptors are released from kinesins, is currently unknown (Figure 1–figure supplement 2A). Recent studies have introduced the new concept that end-binding proteins (EBs) at the plus end of MTs facilitate cargo release (Guedes-Dias et al. 2019; Pawson et al. 2008; Qu et al. 2019). This concept and the possibility that adhesion molecules determine synaptic partners and synapse locations led us to test a mechanism: Adhesion molecules bind MTs via EBs in the vicinity of inhibitory synapses and facilitate cargo release after achieving synapse specificity. This mechanism should contribute significantly to protein accumulation during early development when the distribution of signaling molecules is inadequate.

**Figure 1.**
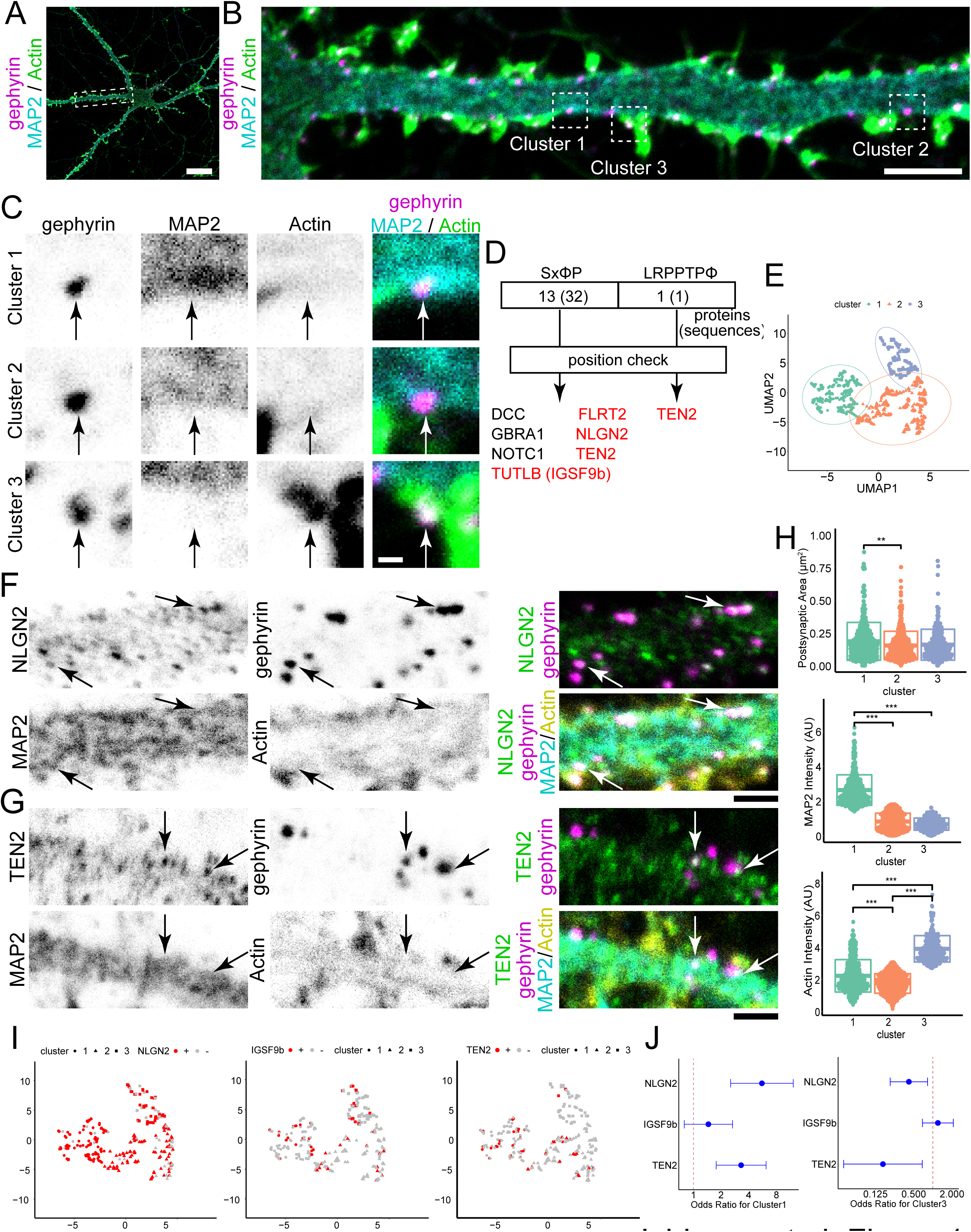
Cluster analysis of inhibitory postsynapses and correlation with TEN2. (A) Image of immunofluorescence staining of gephyrin, MAP2, and actin in DIV20 hippocampal cultured neuron. The dashed box is magnified in (B). Scale bar, 20 μm. (B) Cytoskeletal diversity of inhibitory postsynapses. Cluster 1 is MAP2-rich synapses, cluster 2 is synapses with low levels of both MAP2 and actin, and cluster 3 is actin-rich synapses. Typical synapses are boxed by dash lines with the cluster number attached to each, and an enlarged view is shown in (C). Scale bar, 5 μm. (C) Enlarged view of the synapses belonging to each cluster. Arrows indicate the position of postsynapses. Scale bar, 500 nm. (D) Results of motif search. Each motif was present in 32 and 1 location in each of 13 and 1 protein types. After checking whether these sequences were intracellular or extracellular, the number of candidate proteins was narrowed down to 7. Of these, those belonging to adhesion molecules are shown in red. (E) Plots showing the results of cluster analysis. Inhibitory postsynapses were evaluated by 3-dimensional parameters of synaptic area, MAP2 intensity, and actin intensity. After being reduced to 2 dimensions by UMAP, cluster analysis was performed with the number of clusters pre-specified as 3. The number of synapses belonging to each cluster was 315, 413, and 212 observed by three independent experiments. (F) Images of immunofluorescence staining of Neuroligin-2 (NLGN2), gephyrin, MAP2, and actinin DIV20 hippocampal cultured neuron. Arrows indicate representative NLGN2-positive synapses belonging to cluster 1. Scale bar, 2 μm. (G) Images of immunofluorescence staining of Teneurin-2 (TEN2), gephyrin, MAP2, and actin in DIV20 hippocampal cultured neuron. Arrows indicate representative TEN2-positive synapses belonging to cluster 1. Scale bar, 2 μm. (H) Comparison between clusters for each parameter. Synaptic area: One-way ANOVA showed a significant difference (p = 0.0019), Tukey multiple comparison showed significant difference between cluster 1 and 2 (p = 0.0016). MAP2 intensity: One-way ANOVA showed a significant difference (p < 2e^-16^), Tukey multiple comparison showed significant differences between cluster 1 and 2 (p < 1e^-07^) and between cluster 1 and 3 (p < 1e^-07^). Actin intensity: One-way ANOVA showed a significant difference (p < 2e^-16^), Tukey multiple comparison showed significant differences between cluster 1 and 2 (p < 1e^-07^), between cluster 2 and 3 (p < 1e^-07^) and between cluster 1 and 3 (p < 1e^-07^). Sample size is the same as (E). **p < 0.01, ***p < 0.001. (I) Cluster analysis and the relationship between the positivity and negativity of each adhesion molecule. The calculation results by UMAP are the same as in (E). The number of NLGN2 positive and negative synapses are 228 and 65. The number of IgSF9b positive and negative synapses are 53 and 283. The number of TEN2 positive and negative synapses are 49 and 262. TEN2 positive had very little classification to cluster 3, only 2 synapses. (J) Odds ratio and 95% confidence interval for each adhesion molecule for cluster 1 and 3. For cluster 1: NLGN2, 5.57 (2.54-12.2); IgSF9b, 1.45 (0.80-2.66); TEN2, 3.30 (1.77-6.17). For cluster 3: NLGN2, 0.42 (0.21-0.82); IgSF9b, 1.20 (0.69-2.09); TEN2, 0.16 (0.04-0.68).

To narrow down candidate MT recruiters, we performed a motif search for possible binding to EB1. This search was based on a previous proteomics study that explored the proteins present in the synaptic cleft (Loh et al., 2016). Proteins with two motifs proven to bind to EB1, SxφP, and LxxPTPφ in the intracellular domain were searched for in the results of the proteomic study (Figure 1–figure supplement 2B; Honnappa et al., 2009; Kumar et al., 2017). After confirming the location of the motifs, we narrowed the list of seven proteins as candidate molecules (Figure 1D; Table S1-S6). Neuroligin-2 (NLGN2), immunoglobulin superfamily member 9 B (IgSF9b), and TEN2 were tested among these candidates because of their functions as adhesion molecules and antibody availability (Poulopoulos et al., 2009; Sando et al., 2019; Woo et al., 2013). Next, we performed four-color immunofluorescence staining of each protein along with gephyrin, MAP2, and actin for cluster analysis at DIV 20 (Figures 1E-G, and Figure 1–figure supplement 2D). The clustering results showed that postsynapses were clustered according to whether they were MAP2-rich or actin-rich, as in preliminary experiments (Figure 1H). The postsynaptic area was slightly higher in cluster 1 but had little effect on clustering. Odds ratios were calculated for each cluster for NLGN2, IgSF9b, and TEN2, and the results showed that NLGN2- or TEN2-positive inhibitory postsynapses were more likely to be classified in cluster 1 and not in cluster 3, whereas IgSF9b showed no trend regarding classification (Figures 1I, 1J, and Figure 1–figure supplement 2C). When only MAP2 intensity was compared using classical single-parameter comparison, only NLGN2 and TEN2 showed significant differences between postsynaptic positivity and negativity (Figure 1–figure supplement 2E). These results suggest that NLGN2 and TEN2 tend to be more abundant at MAP2-rich post-synapses and are likely MTs recruiters.

To evaluate whether NLGN2 or TEN2 is more suitable as an MT recruiter, we referred to previous electron microscope (EM) studies. EM studies have suggested that few MTs are observed within 100 nm below the inhibitory postsynapse and are located slightly outside the synapse (Gulley & Reese, 1981; Linsalata et al., 2014). Electron tomography data focusing on GABA receptors have also revealed the absence of MTs just below the synapse (Linsalata et al., 2014; Liu et al., 2020). Among these candidates, NLGN2 is observed near the center of the synapse by EM, which is different from the currently suggested MT localization (Takács et al., 2013; Uchigashima et al., 2016). Therefore, we hypothesized that TEN2 plays a more direct causal role, although NLGN2 is closely associated with MAP2-rich synapses. In this study, we focused on TEN2 to elucidate the mechanism by which MTs are recruited to inhibitory post-synapses.

### Interaction with MTs via EB1 by two motifs in TEN2

The screening results suggest that TEN2 may bind to EB1. EB1 is localized at the MT plus end at the endogenous expression level and is observed as the EB1 comet, but it detaches after the polymerization of MTs is completed (Akhmanova & Steinmetz, 2015). Because the EB1 comet can be seen transiently in living cells and its duration is approximately 10–20 seconds, the EB1 comet is not easy to detect in a fixed cell. However, when large amounts of EB1 are expressed, EB1 localizes to the MT lattice (Skube et al., 2010), making it relatively easy to quantitatively evaluate protein-protein interactions using correlation coefficients (Ichinose et al., 2019). Thus, we first expressed large amounts of EB1 in COS-7 cells and measured its correlation coefficient with TEN2.

TEN2N-L, a chimeric protein consisting of an intracellular domain with two EB1-binding motifs connected to the transmembrane domain of TEN2, was coexpressed with EB1 (Figure 2A). The results showed that TEN2N-L strongly colocalized with EB1-TagRFP and MTs (Figures 2B and 2C). In contrast, TEN2TM, which has only a transmembrane domain, and TEN2N-L2mut, which has an amino acid mutation in the EB1 binding motif, did not colocalize with EB1 (Figures 2B and 2D). Therefore, the intracellular domain of TEN2 is capable of interacting with MTs via EB1. To confirm this interaction, we expressed TEN2N-L and TEN2TM in neurons and observed their colocalization with endogenous EB1 in dendrites. As expected, colocalization of EB1 was observed in a portion of TEN2N-L cells (Figures 2E and 2F). However, uniform colocalization was not observed, as in the experiment with COS-7. This could be attributed to the short EB1 comet dwell time. These experiments suggested that the intracellular domain of TEN2 interacts with MTs via EB1.

**Figure 2.**
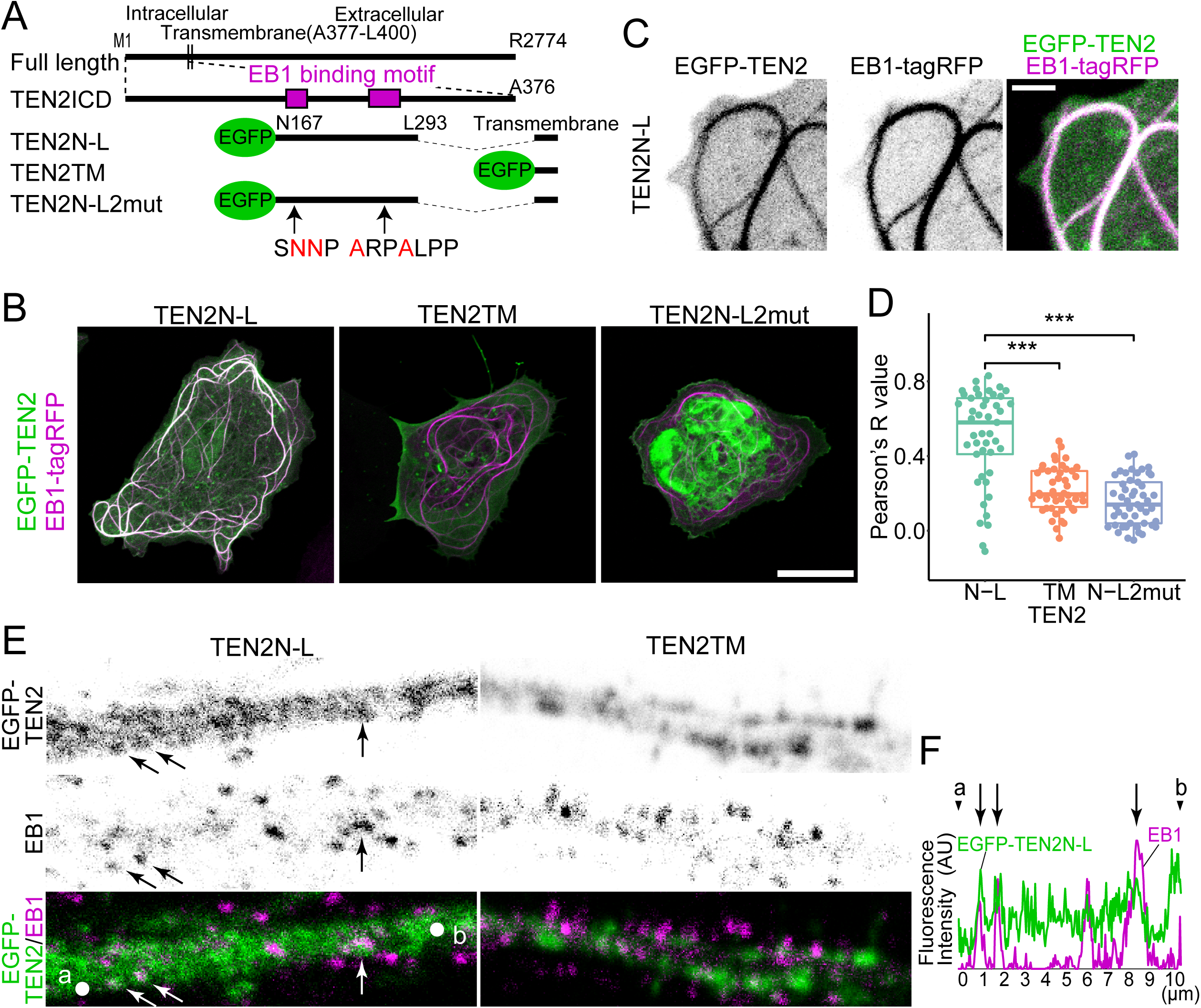
Interaction of two TEN2 motifs with MTs via EB1. (A) Overview of the truncated mutant. TEN2N-L was designed to contain the two EB1 binding motifs detected by motif search. TEN2N-L2mut has amino acid mutations in two EB1 binding motifs. All proteins have transmembrane domains with additional topogenic sequences. (B) Co-expression of each truncated mutant with EB1 in COS-7 cells. Cells with MTs patterns of over-expression of EB1 were observed. TEN2N-L colocalized well with EB1 compared to other mutants, suggesting that TEN2 interacts with EB1. Scale bar indicates 20 μm. (C) Highly magnified image of COS-7 cells expressing TEN2N-L. Scale bar indicates 2 μm. (D) Individual plots and box plots showing the quantitative analysis results of the colocalization index between each TEN2 and EB1, based on correlation coefficients. The median Pearson’s correlation coefficient between TEN2N-L and EB1 was 0.58, which was significantly different from that of TEN2TM (0.195; p = 1.3e^-7^), and TEN2N-L 2mut (0.14; p = 2.9e^-9^) by Pairwise comparisons using Wilcoxon rank sum test after Kruskal-Wallis rank sum test (p = 5.0e^-11^). The total number of cells observed was 46, 46 and 49, respectively. ***p < 0.001. (E) Neurons expressing EGFP-TEN2 fixed in methanol after detergent treatment and immunostained with EB1 and MAP2. Some colocalization of TEN2N-L and EB1 is observed in neurons expressing TEN2N-L (arrows). Note that the EB1 signal in the absence of MAP2 signal suggests that it is EB1 in the axon. Scale bar indicates 2 μm. (F) Line graph showing the signal intensity of EGFP-TEN2 and endogenous EB1. The horizontal axis shows the length, and the vertical axis shows the fluorescence intensity. Points indicated by letters and arrowheads represent positions of a and b in (E). The arrows also correspond to the positions indicated in (E), suggesting some colocalization of TEN2N-L and EB1.

### Enhanced MT-trapping capability by immobilization of TEN2

To investigate whether the interaction between TEN2 and EB1 was related to the recruitment of dynamic MTs, we first observed the arrangement of MTs. The fusion proteins TEN2N-L or TEN2TM, with an HA tag added to the extracellular domain, were expressed in COS-7 cells. Immunofluorescence staining of living cells with anti-HA tag antibodies confirmed that the EB1 binding motif was correctly located in the cytoplasm (Figure 3B). Under these conditions, there was no difference in MT arrangement between TEN2N-L and TEN2TM (Figure 3C, lower panel). TEN2 is endogenously bound to its binding partner at the synapse, immobilized, and restricted from lateral diffusion. Therefore, to mimic the dynamics of endogenous TEN2 at synapses, we immobilized TEN2 using anti-HA tag antibodies on coverslips (Figure 3A). The results showed a region where MTs were absent at the periphery of cells expressing TEN2N-L cultured on coverslips coated with an anti-HA tag antibody (Figures 3C and 3D). At the same time, structures resembling actin protrusions were observed. This resembles retraction fibers (RF). The ratio of cells with RF-like structures increased in TEN2N-L on the anti-HA tag compared to the other conditions (Figure 3E). These results suggest that TEN2N-L affects cytoskeletal structure, and immobilization of TEN2 in the extracellular domain enhances its function.

**Figure 3.**
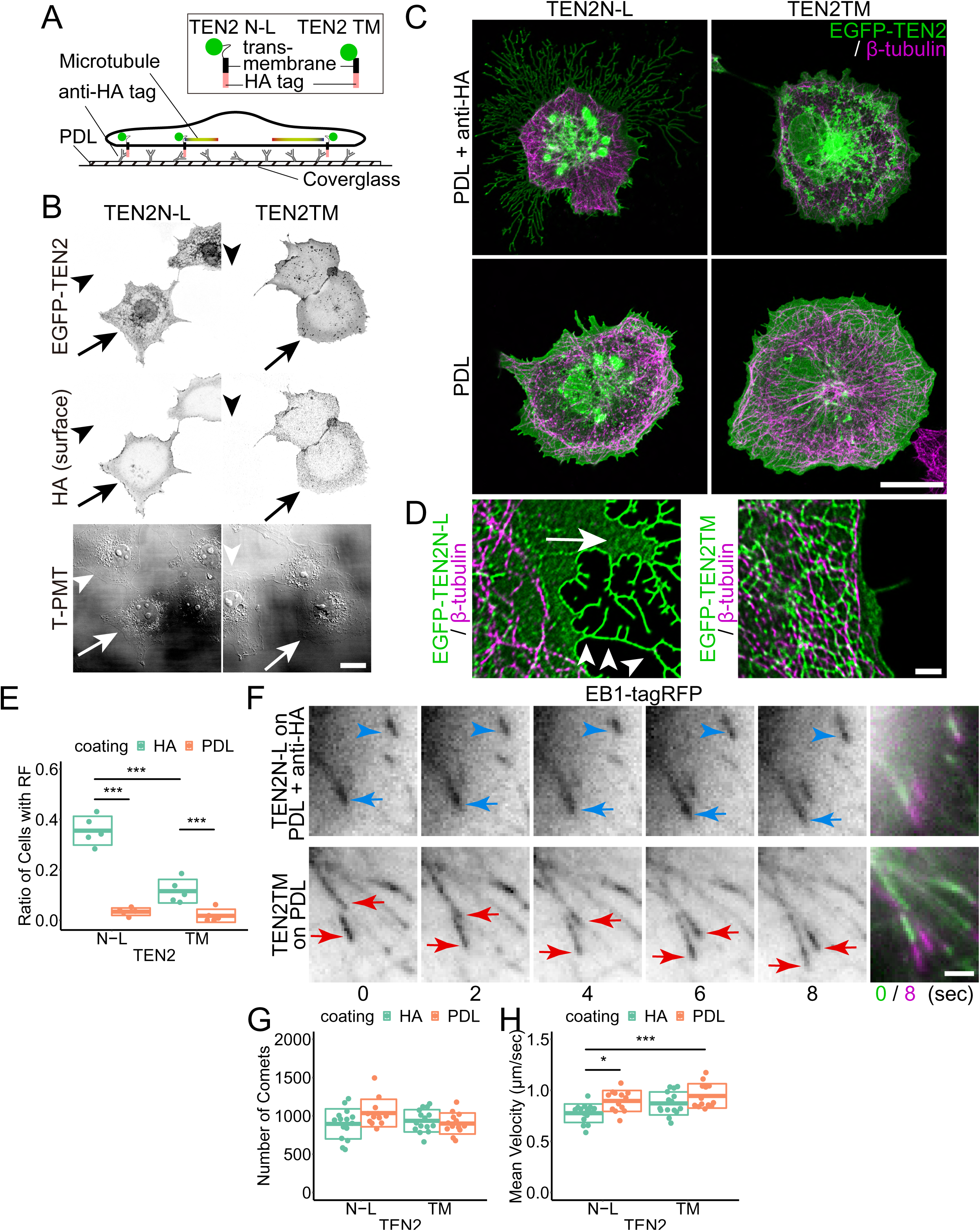
Enhanced MT-trapping capability by immobilization of TEN2. (A) All cover slips were first coated with poly-D-lysine (PDL). Coverslips coated with anti-HA tag antibody and coverslips not coated with antibody were then prepared. Each TEN2 was designed with an additional HA tag in the extracellular domain. The coated antibodies capture these HA tags, and each TEN2 is immobilized. (B) Surface expression of the HA tag was confirmed by expressing TEN2N-L and TEN2TM in COS-7 on a cover slip coated only with PDL. EGFP-TEN2 is also observed on the nuclear membrane and intracellular vesicles, but HA tag does not stain these structures, confirming its presence on the cell membrane (arrows). In non-transfected cells (arrowheads), no EGFP and HA signals were observed, suggesting successful orientation, as outlined in (A). T-PMT shows the signal obtained by the photomultiplier for transmitted light (T-PMT) when the 488-nm laser is illuminated. Scale bar, 20 μm. (C) Confocal microscopic image showing morphological changes of COS-7 cells expressing TEN2N-L, particularly at the cell periphery, when cultured on Anti-HA tag. Scale bar, 20 μm. (D) Super-resolution microscopic images (SRRF-Stream) of cell morphology changes induced by immobilized TEN2N-L. Detailed observation shows MTs-free regions at the cell periphery (arrow) and branching protrusion or retraction fiber -like structures (arrowheads). Scale bar, 20 μm. (E) Plots and crossbars (mean ± SD) showing the results of quantitative analysis of the ratio of COS-7 showing morphological changes in each condition. Each mean ± SD is as follows: TEN2N-L on antibody, 0.36 ± 0.057; TEN2N-L without antibody, 0.034 ± 0.019; TEN2TM on antibody, 0.11 ± 0.047; TEN2TM without antibody, 0.018 ± 0.026. A two-way ANOVA showed a significant difference between TEN2 types (p = 2.10e^-06^) and coating types (p = 2.71e^-09^). Since there was also a significant difference in the interaction (p = 1.07e^-05^), a post-hoc test using the Tukey multiple comparison was performed. n = 5 observations from three independent experiments. The total number of cells observed was 756, 597, 796, and 995, respectively. **p < 0.01, ***p < 0.001. (F) Time-lapse images of EB1 comet. Cells expressing TEN2N-L immobilized by antibodies show slower (blue arrows) or almost immobile (blue arrowheads) EB1 comets, compared to those (red arrows) in cells expressing TEN2TM without immobilization. Scale bar, 2 μm. (G) The number of comets for each observation ranged from 583–1495. mean ± SD was 895 ± 196, 1028 ± 175, 900 ± 137 and 935 ± 137, respectively. There was no significant difference between TEN2 types (p = 0.40) and coating types (p = 0.28) by two-way ANOVA. n = 14, 12, 12 and 14 observations, respectively, from three independent experiments. n.s., not significant. (H) The mean velocities of the detected comets. mean ± SD was 0.78 ± 0.09, 0.90 ± 0.10, 0.94 ± 0.19 and 0.87 ± 0.11 (μm/sec), respectively. A two-way ANOVA showed a significant difference between TEN2 types (p = 0.011) and coating types (p = 0.0012). Since there was also a significant difference in the interaction (p = 0.011), a post-hoc test using the Tukey multiple comparison was performed (p = 0.022 between TEN2N-L on HA antibody and TEN2N-L on PDL; p = 5.8 e^-05^ between TEN2N-L on HA antibody and TEN2TM on PDL). Observation number is same as (G). *p < 0.05, ***p < 0.001.

Since immobilization affects cell morphology and the cytoskeleton, the interaction between TEN2 and EB1 may also be altered by TEN2 immobilization. To test this, we performed live imaging of dynamic MTs by observing EB1-tagRFP in cells immobilized with TEN2. Since EB1 overexpression itself may affect MT dynamics (Skube et al., 2010), we attempted to minimize the expression levels as much as possible. EB1-tagRFP and each TEN2 truncated mutant were co-transfected into COS-7 cells cultured on coverslips coated with poly-D-lysine (PDL) or anti-HA tag antibodies. Observations were made using a TIRF microscope so that EB1 analysis could be restricted just beneath the plasma membrane. As a result of imaging for 100 seconds at 2-second intervals, comet mobility was slower in TEN2N-L-transfected cells, but stable and immobile comets were also observed (Figure 3F; Figure 3–videos 1 and 2).

The number of comets in each observation was not significantly different (Figure 3G), suggesting no difference in the frequency of comet formation despite variations in cell size, morphology, and transfection volume. Next, we quantified the comet velocity and found that the velocity was significantly reduced in cells with immobilized TEN2N-L on anti-HA tag antibody coating (Figure 3H). These results suggest that TEN2 traps dynamic MTs by interacting with EB1, and that this function is enhanced by immobilization.

### TEN2 localization at the semi-periphery region of the inhibitory postsynapse

TEN2 is a transmembrane protein in synapses. However, the precise localization of TEN2-containing molecular adhesion complexes remains controversial. Some reports suggest that the complex resides in the presynaptic membrane, whereas others suggest that it resides in the postsynaptic membrane (Mosca et al., 2012; O’Sullivan et al., 2012; Sando et al., 2019; Silva et al., 2011; Vysokov et al., 2018). To prevent incorrect identification due to differences in antibodies, we first observed TEN2 localization using dissociated hippocampal cultures prepared from TEN2 knock-in (KI) mice. Mice with a 3xHA tag inserted immediately before the STOP codon of TEN2 in exon 29 were generated using the CRISPR/Cas9 system (Figure 4–figure supplement 1A-D).

Primary cultured hippocampal neurons at DIV 14-15 were co-stained with antibodies against the HA tag and gephyrin and observed with a confocal microscope (Figures 4A and 4B). The results showed that TEN2 and gephyrin colocalized, and the rate of colocalization was consistent with the results of immunofluorescence staining with antibodies against the intrinsic intracellular domain (1E). These results suggest that all antibodies and KI mice can withstand proper analysis.

**Figure 4.**
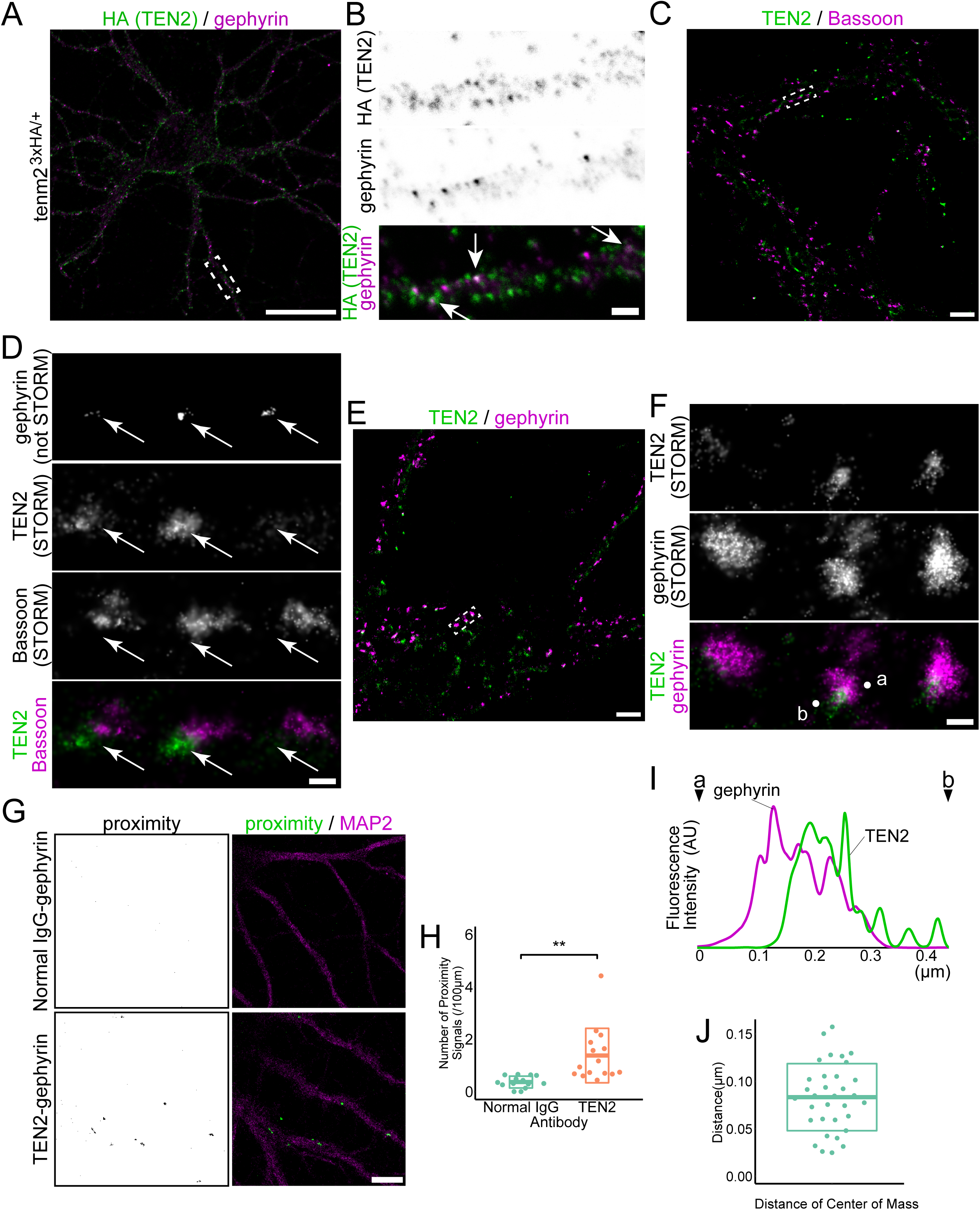
TEN2 localization at the semi-periphery region of the inhibitory postsynapse. (A) Image of immunofluorescence staining of HA tag and gephyrin in knock-in neuron. The dashed box is magnified in (B). Scale bar, 50 μm. (B) Colocalization of gephyrin and HA tag signals was observed, suggesting that TEN2 is also localized at inhibitory synapses. Arrows represent colocalization. Scale bars indicate 2 μm. (C) dSTORM Images of TEN2 immunostained with anti-ICD antibody and bassoon. The dashed box is magnified in (D). Scale bar, 2 μm. (D) Magnified images. Colocalization of TEN2 and bassoon signals was not obvious, suggesting that TEN2 is more localized at postsynapse than presynapse. Gephyrin is labeled with AF488, but STORM images cannot be constructed under the conditions. Thus, arrows just represent that these synapses are inhibitory, not actual gephyrin position. Scale bar, 200 nm. (E) dSTORM Images of TEN2 and gephyrin. The dashed box is magnified in (F). Scale bar, 2 μm. (F) Colocalization of TEN2 and gephyrin. The majority of TEN2 overlaps with gephyrin, but their centers of mass do not necessarily coincide. Scale bar, 200 nm. (G) Images showing the results of the proximity ligation assay. When the proximity ligation assay was performed using antibodies against TEN2 and gephyrin, a signal indicating proximity of less than 20 nm could be detected. On the other hand, no signal was obtained in the negative control. Scale bar, 10 μm. (H) The number of proximity signals per 100 μm. mean ± SD was 0.37 ± 0.23 and 1.38 ± 1.04, respectively. Welch’s t-test showed a significant difference between negative control and TEN2 with respect to proximity to gephyrin (p = 0.0021). The number of observed neurons is 14 and 15, respectively, from three independent experiments. **p < 0.01. (I) Line graph showing the signal intensity of TEN2 and gephyrin. The horizontal axis shows the length, and the vertical axis shows the fluorescence intensity. Points indicated by letters and arrowheads represent positions of a and b in (F). (J) Distance between the centers of mass of TEN2 and gephyrin when observed in dSTORM. The mean ± SD was 83.3 ± 35.3. The number of synapses observed was 33, from three independent experiments.

Next, we performed antigen-antibody reactions on live neurons before fixation and found that a portion of TEN2 was located on the dendritic shaft (Figure 4–figure supplement 1F and G). These results suggest that TEN2 localizes and functions in the plasma membrane of dendrites. On the other hand, presynaptic TEN2 has been reported to bind to latrophilin in the postsynaptic area and induce differentiation of excitatory postsynapses (Sando et al., 2019; Sando & Südhof, 2021). The results of co-staining with actin and PSD-95 indicated that TEN2 was also localized at the excitatory synapse (Figure 4–figure supplement 1H-K). Thus, our results suggest that TEN2 is expressed and functions in the plasma membrane at both excitatory and inhibitory synapses.

However, it is impossible to detect whether TEN2 is located at the presynaptic or postsynaptic membrane because of limitations in the resolution of the microscope. Therefore, we next observed precise localization using one of super-resolution microscope (SRM), stochastic optical reconstruction microscope (STORM). First, to determine whether TEN2 is present in the presynapse or postsynapse at inhibitory synapses, we co-stained the cells with an antibody against the intracellular domain of TEN2 (anti-ICD) and bassoon, a marker of presynapse, at DIV 21. The STORM image shows little overlap between TEN2 and the bassoon (Figures 4C and 4E). It should be noted that this result alone does not rule out a presynaptic presence, since the size of the antibody is almost the same (∼20 nm) as that of the synaptic cleft. Thus, these results suggest that TEN2 is more abundant at postsynapses than at presynapses of inhibitory synapses. It should also be noted that the combination of available lasers, dyes, and reducing agents limits the reliability of the localization at the super-resolution level to two molecules in our experimental system. TEN2 was labeled with CF568, bassoon with Alexa Fluor (AF) 647, and gephyrin with AF488. AF488 recordings only indicated that these synapses were inhibitory and could not show precise localization. To observe the exact localization of TEN2 and gephyrin, different combinations of dyes were used in the STORM. An overlap between TEN2 and gephyrin was observed, suggesting that TEN2 was present at the inhibitory synapses (Figures 4D and 4F). A proximity-ligation assay was performed to confirm this result (Söderberg et al., 2006). In this assay, two antibodies were immunostained, and if they were close (∼ 20 nm), oligonucleotides fused to the antibodies were ligated to produce circular DNA. Rolling circle amplification was performed to detect proximity by incorporating dyes into the dNTPs. The results showed that the proximity between TEN2 and gephyrin was significantly greater than that between normal IgG and gephyrin, which was used as the negative control (Figures 4G and 4H). This result supports our STORM data and suggests that the effect of signal misalignment between channels owing to drift and other factors is minimal. Interestingly, the puncta of TEN2 and gephyrin were not always perfectly colocalization. Therefore, we measured the distance between the centers of mass of the fluorescence intensities of each punctum and found that they were 85 nm apart (Figures 4F, 4I, and 4J). Because this super-resolution observation was performed in 2D-STORM, the actual distance would be slightly longer. Considering the width of the inhibitory postsynapse (approximately 500 nm), this distance is not as far outward as the perisynapse. These results suggest that TEN2 is more abundant at inhibitory postsynapses and is present in semi-peripheral regions.

### Inhibitory postsynapse maturation induced by postsynaptic TEN2

TEN2 overexpression in non-neuronal cells induces maturation of both excitatory and inhibitory synapses (Sando et al., 2019). To investigate whether TEN2 also induces synaptic maturation in neurons, we knocked down TEN2 in primary hippocampal cultures using RNAi. Knockdown was performed using a vector-based shRNA (Figure 5–figure supplement 1A). The vector has an shRNA sequence downstream of the mouse U6 promoter and a TagRFP sequence downstream of the SV40 promoter, allowing TagRFP to be expressed as a volume marker. The half-life of TEN2 from DIV11 in primary cultured rat hippocampal neurons is 1.42 days (Heo et al., 2018). Based on these data, we transfected DIV11 cells with TEN2 knockdown vectors and fixed and immunostained them three days after transfection.

TEN2 expression levels in the cell body were quantified by immunofluorescence staining with anti-TEN2 to confirm knockdown (Figure 5–figure supplement 1B and C). In neurons transfected with the TEN2 knockdown vector, the mean signal intensity of TEN2 was significantly reduced to 52% compared to neurons transfected with the control vector (Figure 5–figure supplement 1D). Under these conditions, we found that the number of gephyrin puncta was significantly reduced in neurons transfected with the TEN2 knockdown vector (Figures 5A and 5B), suggesting that postsynaptic TEN2 was involved in the maturation of inhibitory postsynapses.

**Figure 5.**
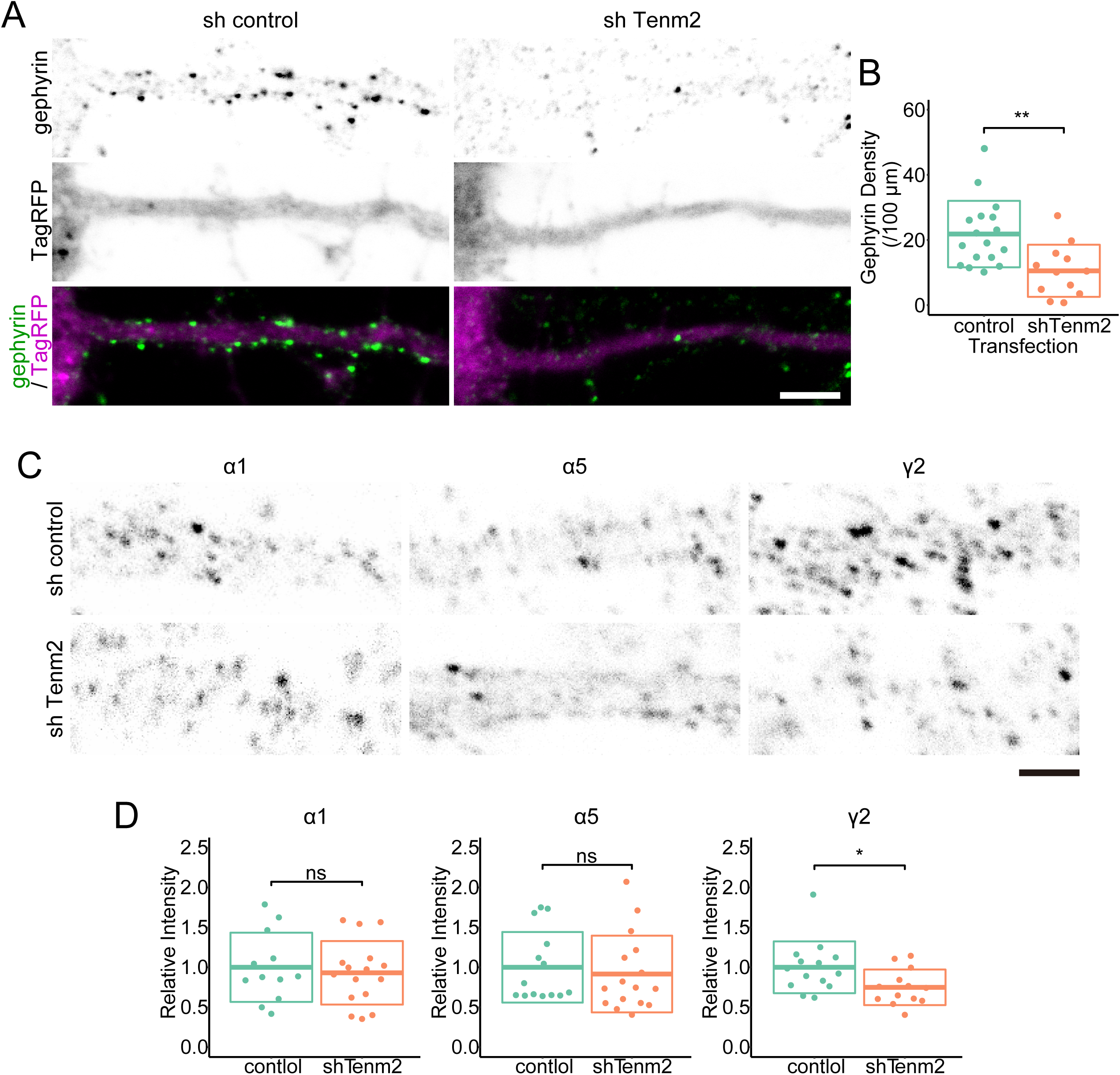
Inhibitory postsynapse maturation induced by postsynaptic TEN2. (A) Magnified images of knockdown neurons and immunofluorescence staining of gephyrin. Gephyrin accumulation was reduced in TEN2 knockdown neurons. The scale bar is 5 μm. (B) Plots and crossbars (mean ± SD) quantifying the density of gephyrin puncta per 100 μm. It was 21.8 ± 10.2 in the neurons transfected with the control vector, and 10.5 ± 8.0 in neurons transfected with TEN2 knockdown vector (p = 0.0025). n = 16 control neurons and 12 knockdown neurons, from four independent experiments. **p < 0.01. (C) Magnified images of knockdown neurons and immunofluorescence staining of GABAA receptors subunit α1, α5 and γ2. Of these subunits, only the γ2 receptor is downregulated in TEN2 knockdown neurons. The scale bar is 2 μm. (D) Plots and cross bars (mean ± SD) quantifying relative fluorescence intensity of GABAA receptor subunits. The fluorescence intensities of receptors present in dendrites within 100 µm from the cell body were quantified comparatively. mean ± SD were 1 ± 0.43 and 0.93 ± 0.40 for α1, 1 ± 0.40 and 0.92 ± 0.48 for α5, and 1 ± 0.32 and 0.75 ± 0.22 for γ2. Results of Welch’s t-test showed that α1 (p = 0.67) and α5 (p = 0.62) were not significantly different between control and TEN2 knockdown cells, but γ2 (p = 0.027) was predominantly reduced in TEN2 knockdown cells. The number of cells observed were 12, 16, 14, 16, 14, and 13, respectively from three independent experiments. *p < 0.05.

Although we used gephyrin as a marker of inhibitory synapses throughout this study, we examined how TEN2 knockdown affects the localization of GABA_A_ receptors. There are 19 GABA_A_ receptor subunits. Of these, the GABA_A_ receptor localized at inhibitory synapses forms a hetero-pentamer consisting of two α1-3 subunits, two β1-3 subunits, and one γ2 subunit, in the order γ2-β-α-β-α, counterclockwise from the extracellular view. In TEN2 knockdown neurons, The GABA_A_ receptor subunits α1, α5, and γ2 were quantified. We found that only γ2 expression was significantly reduced by TEN2 knockdown (Figures 5C and 5D). Since γ2 is present in all synaptic GABA_A_ receptors, a significant decrease in gephyrin would lead to a decrease in γ2. However, α1 is a subunit present only in a specific subset of synaptic GABA_A_ receptors, so TEN2 may not have a significant effect on synaptic maturation in this subset. α5 is a subunit of extrasynaptic GABA_A_ receptors and is not integrated into intrasynaptic GABA_A_ receptors. Therefore, it seems quite natural that TEN2, which functions via adhesion at inhibitory synapses, does not affect the distribution of extracellular GABA_A_ receptors. These results suggest that TEN2 is involved in postsynaptic maturation at inhibitory synapses and promotes accumulation of gephyrin and GABA_A_ receptors at these synapses.

### TEN2-MT interactions lead to maturation of the inhibitory postsynapse

As shown thus far, the interaction between TEN2N-L and MTs is constitutively active. However, TEN2N-L cannot bind to its partners in the synaptic cleft because of the loss of its extracellular domain. Therefore, we assumed that we could inhibit the synaptogenic function of endogenous TEN2 in a dominant-negative (DN) manner by immobilizing TEN2N-L throughout the dendrites. First, we evaluated the effects of non-immobilized TEN2 on primary cultured hippocampal neurons. Gephyrin accumulation was affected even without TEN2 immobilization (Figures 6A and 6B). Furthermore, when MAP2 was considered as MTs, MTs in TEN2N-L-expressing dendrites were sparser than those in TEN2TM and biased toward both ends parallel to the axial direction (Figures 6A and 6C). The ratio of neurons with this MTs distribution significantly increased in TEN2N-L-expressing neurons (Figure 6D).

**Figure 6.**
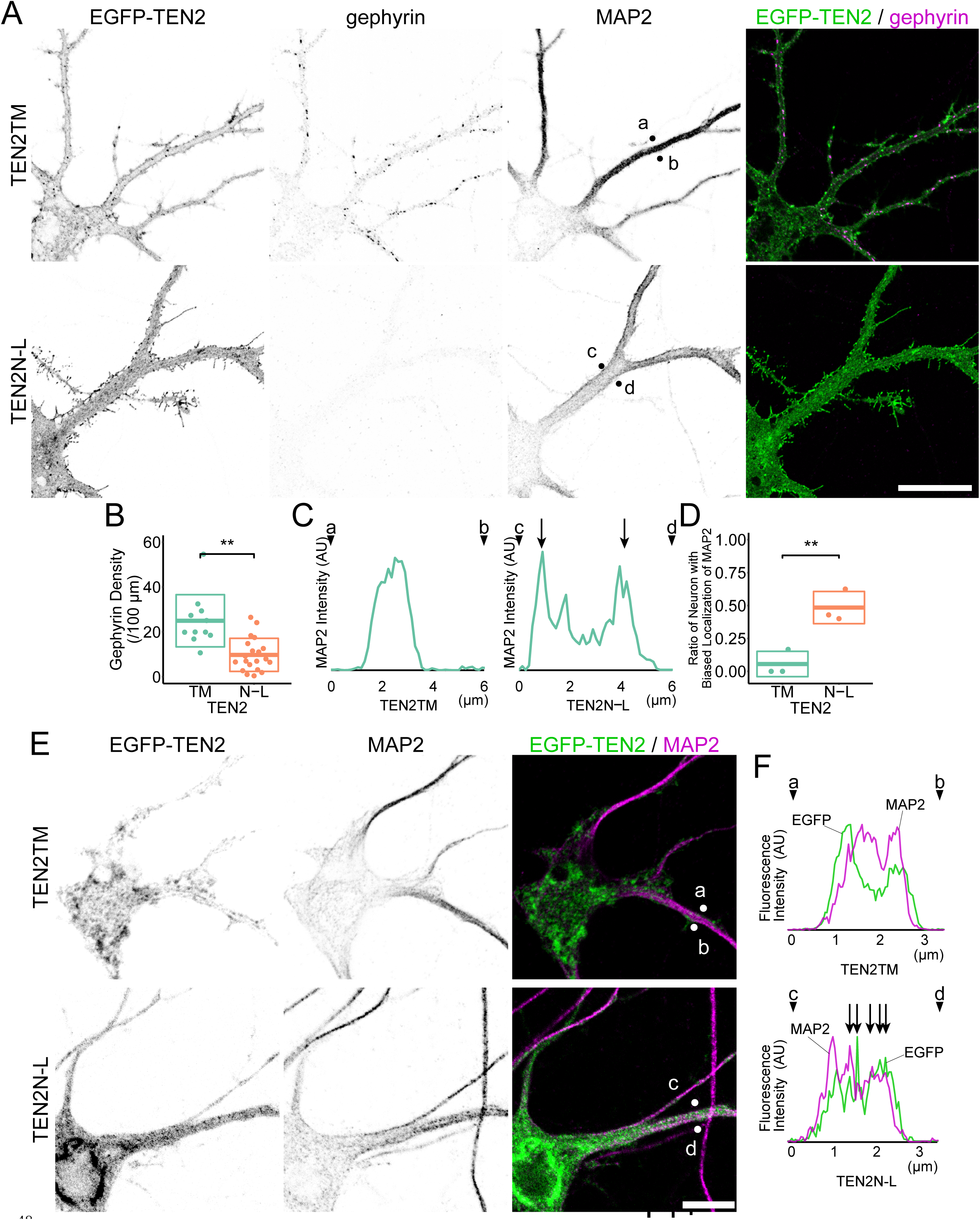
TEN2-MT interactions lead to maturation of the inhibitory synapse. (A) Confocal imaging of gephyrin accumulation and MAP2 in neurons expressing each TEN2. Higher gephyrin accumulation was observed in control (TEN2TM) neurons, whereas it was reduced in dominant-negative TEN2N-L. In addition, biased MAP2 were observed in the TEN2N-L. Scale bar, 20 μm. (B) Plots and crossbars (mean ± SD) showing densities of gephyrin puncta per 100 μm. It was 26.2 ± 15.5 in neurons with TEN2TM and 13.3 ± 8.29 in TEN2N-L (p = 0.0014). n = 11 control neurons and 20 DN from three independent experiments. **p < 0.01. (C) Line graph showing the signal intensity of MAP2. The horizontal axis shows the length, and the vertical axis shows the fluorescence intensity. Points indicated by letters and arrowheads represent positions of a-d in (A). In control neurons, the MAP2 signal is strongly observed around the dendrite axis, suggesting that MTs are strongly bundled. In DNs, on the other hand, the MAP2 peak is biased to be located just below the membrane in the direction parallel to the axis and is sparse near the axis. This suggests that DN on the membrane recruits MTs. (D) Ratio of neurons with membrane-biased MTs. mean ± SD were 0.056 ± 0.096 and 0.48 ± 0.12, which were significantly different (p = 0.01) by Welch’s t-test. Observations were based on three independent trials. **p < 0.01. (E) TEN2N-L interacting with the cytoskeleton. Living neurons were treated with detergent to remove proteins that do not interact with the cytoskeleton partially and then quickly fixed with ice-cold methanol. TEN2TM was partially eluted by detergent treatment. In comparison, TEN2N-L was eluted less and mainly colocalized with MAP2 as a dendritic MTs marker. Scale bar, 10 μm. (F) Line graph showing the signal intensity of EGFP-TEN2 and MAP2. The horizontal axis shows the length, and the vertical axis shows the fluorescence intensity. Points indicated by letters and arrowheads represent positions of a-d in (E). In the control neurons, MAP2 signal is inside the EGFP signal suggesting that these proteins do not colocalize. Meanwhile, in DN, the positions of the peaks of TEN2N-L and MAP2 overlap (arrows), suggesting that these proteins colocalize.

To determine whether TEN2N-L interacts with neuronal MTs, we extracted membrane proteins that did not interact with the cytoskeleton using saponins prior to methanol fixation. EGFP signals were sparsely detected in control neurons, and no colocalization with MAP2 was observed. On the other hand, in TEN2N-L-expressing neurons, dense EGFP signals remained and colocalization with MAP2 was observed, which was biased at both ends parallel to the axial direction (Figure 6E and 6F). These results suggest that TEN2N-L indiscriminately interacts with MTs throughout dendrites, thereby disrupting the endogenous TEN2-MT interaction and synaptic maturation. This suggests that endogenous TEN2-MTs interaction promotes the maturation of inhibitory synapses.

## Discussion

We found that inhibitory postsynapses could be categorized into three clusters based on the amount of MAP2 and actin: MAP2-rich postsynapses, actin-rich postsynapses, and postsynapses with low levels of both MAP2 and actin. Because TEN2, which has EB1 binding motif, tends to localize MAP2-rich postsynapses, we analyzed the relationship between TEN2 and MTs. We found that TEN2 traps MTs via EB1, and that this function is enhanced by TEN2 immobilization. TEN2 was localized to the postsynaptic semi-periphery region, and knockdown of TEN2 suppressed gephyrin clustering. Furthermore, expression of the dominant-negative form also suppressed gephyrin clustering. Considering these results and those of previous studies, we propose the following working model to explain the function of TEN2 in inhibitory synaptic maturation (Figure 7). In wild-type neurons, TEN2 localizes to the postsynaptic semi-periphery region and is anchored by a presynaptic binding partner; the interaction of TEN2 with dynamic MTs provides an unloading zone for motor proteins and allows proper exocytosis of synaptic components. In contrast, in neurons expressing the dominant-negative form, binding to MTs is constitutively activated throughout the submembrane, resulting in universal interaction between MTs and the submembrane. Therefore, motor proteins cannot select their destination and the components do not accumulate. In knockdown neurons, motor proteins cannot select a destination because the interaction between MTs and the submembrane does not occur.

**Figure 7.**
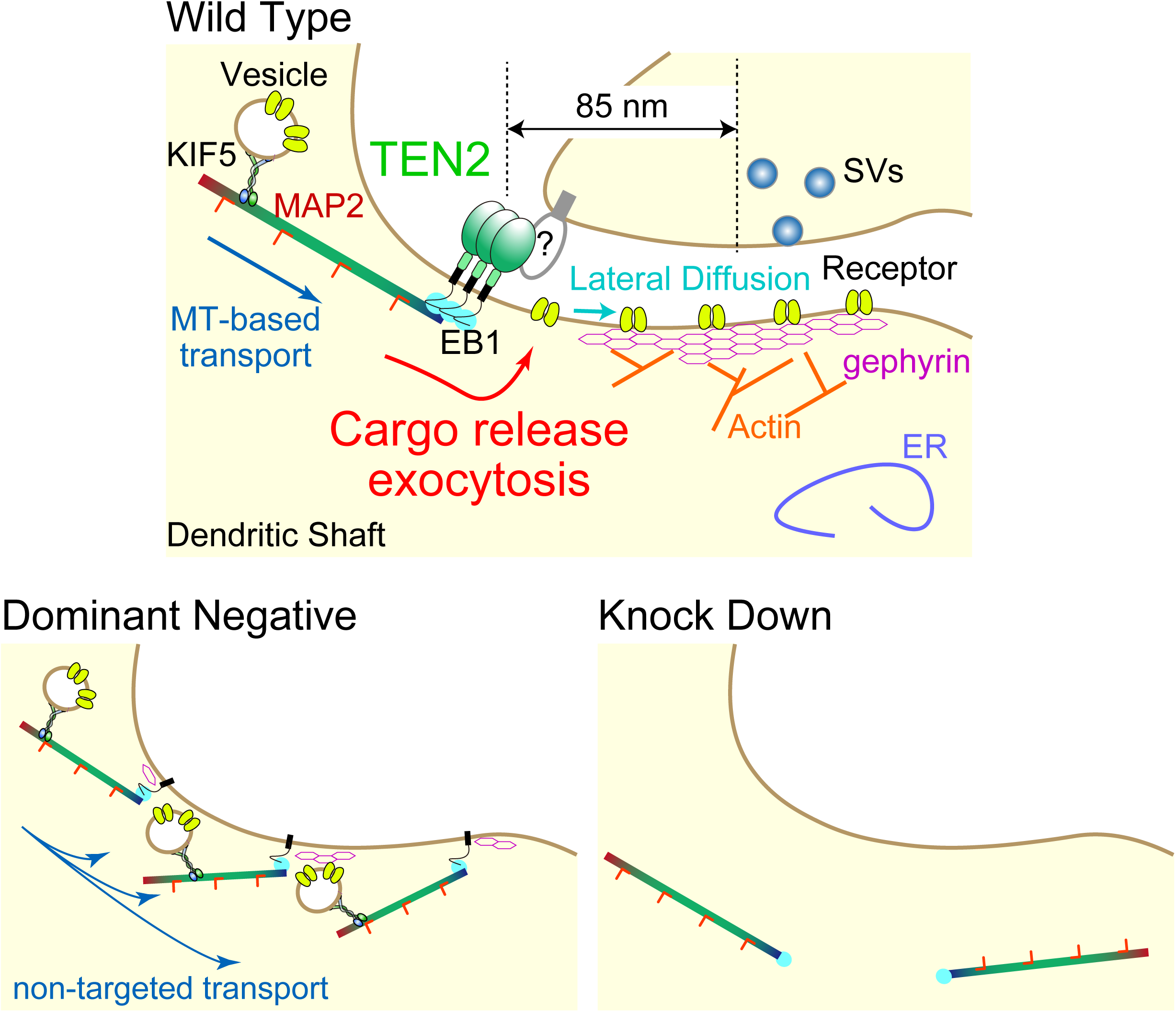
Teneurin-2 at the synapse construction site is a signpost for cargo unloading from motor proteins. Working model derived from this study. Wild-type: interaction of TEN2 with dynamic MTs provides an unloading zone for motor proteins, allowing proper transport of synapse components in the semi-peripheral region of the inhibitory postsynapse. Dominant-negative: TEN2 binding to MTs is constitutively activated throughout the dendrite; therefore, the motor protein cannot select a destination. Knockdown neurons: there is no interaction between MTs and the submembrane; hence, the motor protein cannot select a destination.

We conclude that the interaction of TEN2 with MTs is critical for efficient transport of synaptic components to develop inhibitory synapses.

### Role of TEN2-MT interactions in the inhibitory postsynapse

Inhibitory synapses can be categorized into three clusters based on the amount of MAP2 and actin. TEN2, which has an EB1 binding motif, tends to localize to MAP2-rich postsynapses, and we analyzed the relationship between TEN2 and MTs. TEN2 was observed at the semi-periphery of the inhibitory postsynapses; the centers of mass of the fluorescence intensities of TEN2 and gephyrin were 85 nm apart. However, this is based on 2D observations; therefore, the actual distance may be slightly longer, roughly estimated to be ∼120 nm. Considering that the width of postsynapses is generally 500 nm to 1 µm, we determined that they are present in the semi-periphery. This distance would be adequate because if it were located outside the postsynaptic area, it would lose its connection to its presynaptic counterpart, resulting in a reduced MTs trapping function. Indeed, dynein, a retrograde motor protein, has also been observed in this region by EM, which supports our results (Fuhrmann et al., 2002).

The dominant negative form of TEN2 induces abnormalities in MTs arrangement in both COS-7 cells and neurons. These results are consistent with previous experiments showing that in mutants of ten-a, an ortholog of Drosophila teneurin, synaptic MTs are not properly positioned in the neuron, resulting in defects in synaptic transmission (Mosca et al., 2012). Immobilization of TEN2 enhanced its function on the cytoskeleton: In experiments with COS-7, RF-like structures were observed, but no lamellipodia, which should form during cell migration, were found on the opposite side. This may be due to the use of antigen-antibody reactions that do not cause adhesion turnover, such as integrins. Alternatively, the formation of RF-like structures could be interpreted as spontaneous repulsion randomly generated by truncated mutants with inadequate intracellular functions. It has only been discovered that TEN2 is involved in cell repulsion, in addition to cell migration and cell adhesion, and the mechanism is not yet fully understood (Del Toro et al., 2020). Our experimental results are only a prelude to the elucidation of this mechanism, but it is certain that the immobilization of TEN2 causes MTs traps.

The TEN2-MTs interaction is suggested to be crucial not only for cytoskeletal changes but also for cargo distribution. The major regulatory mechanisms of cargo distribution include phosphorylation of kinesins, post-translational modifications of MTs (MTs code), and changes in kinetics between kinesins and MTs by MAPs (MAPs code; Aiken & Holzbaur, 2021; Monroy et al., 2020). Among these, regulation by phosphorylation is disadvantageous for synaptogenesis. This is because protein kinases and phosphatases each have their own distribution patterns; therefore, how to distribute cargo goes back to the question of how to distribute kinases (Ichinose et al., 2015; Ichinose et al., 2019). In other words, once synaptic specificity is converted to a distribution of kinases, there are additional steps for continuity to synaptogenesis. On the other hand, regulatory mechanisms such as MTs state and kinetic changes by MAPs can be controlled only by the interaction between adhesion molecules and MTs, which would allow smooth continuity from synaptic specificity to synaptogenesis. Thus, cargo distribution by “MTs code” and “MAPs code” is more efficient than the kinase theory. However, there are still many unknowns with “MTs code” and “MAPs code”. It has been reported that EB1-kinesin competition reduces the affinity between kinesin and MTs and cargo release is achieved (Guedes-Dias et al., 2019; Qu et al., 2019). This may be favorable for KIF1A, which transports synaptic vesicles in axons, as GTP MTs and EB1 are abundant near the presynapse. However, this model cannot be applied because KIF5, which carries GABA_A_ and glycine receptors, prefers GTP MTs to GDP MTs (Nakata et al., 2011). However, it seems feasible to apply the competition between motors and EBs to the shedding of KIF5 from MTs. The MTs state and MAPs localization of inhibitory post-synapses are unknown at the nanoscale level, and elucidating this is the next challenge. In addition, the interaction between TEN2 and cargo should also be considered in the future because the function of most of the intracellular domains of TEN2, which contain more than 360 amino acids, remains a mystery. Adapter protein HAP1 not only links GABA_A_Rs to KIF5, but also regulates the motility of KIF5 (Twelvetrees et al., 2019). If TEN2 acts on HAP1, it may reduce KIF5 motility and release the cargo.

The TEN2-MTs interaction may also function as a starting point for dynein transport. It has already been reported that dynein transports gephyrin toward the minus end of MTs (Maas et al., 2006). In the present study, we found that the expression of DN-TEN2 in whole dendrites reduced gephyrin accumulation. Because the DN form recruits MTs to form endpoints of anterograde transport throughout the dendrite, receptor transport itself is dominantly activated. At the same time, the origin of retrograde transport is also increased; therefore, our results can be interpreted as an apparent absence of protein accumulation. Thus, one of our future tasks is to elucidate whether adhesion molecules regulate the balance between anterograde and retrograde transport.

Although the TEN2-MTs interaction has been confirmed to promote synaptogenesis, other candidate molecules still need to be investigated. It has been suggested that NLGN2 also localizes to the MAP2-rich synapses. We excluded NLGN2 because it is abundant in the center of inhibitory postsynapses, where few MTs have been observed using EM (Gulley & Reese, 1981; Linsalata et al., 2014). However, postsynapses do not have a size of 500 nm from an early developmental stage. While postsynapses are small, MTs may also be present immediately below the synapse. We also excluded IgSF9b because no correlation was observed in the cluster analysis. This may be related to developmental stage. Previous studies have analyzed IgSF9b using primary cultured neurons at 10-20 DIV, indicating that a substantial number of post-synapses contain IgSF9b at this stage (Woo et al., 2013). However, in the DIV 20 neurons we observed, colocalization of gephyrin and IgSF9b was reduced, suggesting that IgSF9b may be necessary for postsynaptic development, but may not have a function in mature neurons, such as modulating plasticity (Figure 1–figure supplement 2C and D). In the future, the contribution of MTs to inhibitory synapses should be evaluated according to the developmental stages of synapses.

### Binding patterns of TEN2

To investigate the localization of TEN2, a new antibody was generated against its intracellular domain. In addition, HA-tag knock-in mice were generated, and hippocampal neurons from these mice were validated in primary culture; TEN2 was present at both excitatory and inhibitory synapses, and SRM observations showed that TEN2 was abundant at inhibitory postsynapses. Furthermore, gephyrin puncta were reduced in the dendrites of TEN2-knockdown neurons, suggesting that TEN2 plays a role in the inhibitory synapses. These experimental results appear somewhat puzzling, considering previous studies that examined presynaptic and postsynaptic differentiation by expressing TEN2 in non-neuronal cells and co-culturing them with neuronal cultures (Li et al., 2018; Sando et al., 2019). They reported that overexpression of TEN2(SS+) in non-neuronal cells resulted in the detection of GABA_A_ receptor α2 and γ2 subunits in neurons but not vGAT. These results suggested that TEN2 is present in the presynaptic region. However, this method only shows the function of TEN2 binding partners, not TEN2 itself. Thus, our results show that TEN2 functions at inhibitory post-synapses are not incompatible. The simplest hypothesis is that TEN2 binds homophilically at inhibitory synapses but that TEN2 itself does not have a presynaptic differentiation function and requires the help of other molecules.

Indeed, homophilic binding of members of the teneurin family, including TEN2, has been reported in other studies (Beckmann et al., 2013; Berns et al., 2018). However, the lack of homophilic binding of TEN2 in cell aggregation assays refutes this hypothesis (Li et al., 2018). In contrast, another study reported successful cell aggregation assays (Rubin et al., 2002). This study mentions the inhibition of aggregation by the intracellular domain of TEN2, and homophilic binding has been observed only in aggregation assays using only the extracellular domain. It is also interesting to note that conflicting functions of heterologous TEN2-FLRT-latrophilin “binding” at the synapse and “repulsion” within the cell body have been reported (Del Toro et al., 2020). Such conflicting functions are not limited to TEN2, and it has become a common paradigm in recent years that adhesion molecules have both adhesion and repulsion functions, and should therefore be called recognition molecules rather than adhesion molecules (Sanes & Zipursky, 2020). Aggregation assays are often performed by overexpressing recognition molecules in non-neuronal cells. Since recognition molecules can both adhere and repel, if downstream signaling is not sufficiently manipulated, spontaneous repulsion can occur despite intrinsic adhesion, inhibiting cell aggregation and making it difficult to accurately identify the binding partner. Herein lies the limitations of the aggregation assay. This change in cell morphology due to spontaneous repulsion was also observed in our study using COS-7 cells (Figure 3C).

SRM observations revealed that TEN2 localizes both pre- and postsynaptically but in a rather biased postsynaptic proportion (Figure 4G). Given that SRM resolution and antibody size are comparable to the pre-post distance of the synaptic gap (20 nm), SRM cannot be used to assess trans-homophilic binding. Currently, there is insufficient evidence to confirm or deny the trans-homophilic binding. If TEN2 can bind both homophilically and heterophilically, then the bias in the postsynaptic abundance of TEN2 suggests not only trans-homophilic binding, but also a mixture of homophilic and heterophilic binding. It is important for partner choice and synaptogenesis to examine how these two types of binding patterns are functionally used in different ways.

In conclusion, we propose that TEN2-MTs interactions in the cytoplasm serve as the starting and ending points for MTs MT-dependent transport. This mechanism enhances the development of the inhibitory synapses.

## Materials and Methods

### Cell culture

COS-7 (RCB Cat# RCB0539, RRID:CVCL_0224) cells were obtained from the RIKEN Cell Bank and maintained in High-Glucose DMEM (Wako) supplemented with 10% fetal bovine serum (BioWest) in T75 flasks (Thermo Fisher Scientific), not to exceed 80% confluency. Hippocampi were dissected from the brains of ICR mice (Charles River; IMSR Cat# CRL:022, RRID:IMSR_CRL:022) or knock-in mice on embryonic day 16 (E16). No gender determination is done and three or more embryos are used. They were digested with 0.25% trypsin (Thermo Fisher Scientific) in HBSS (Wako) for 15 minutes at 37°C. Dissociated hippocampal cells were plated at 2 × 10^4^ cells per well on Lab-Tek II 8-well chamber coverglasses (Thermo Fisher Scientific) coated with polyethylenimine and BioCoat poly-D-lysine (Corning). All primary cells were cultured in MEM (Thermo Fisher Scientific) supplemented with 1 mM pyruvate (Thermo Fisher Scientific), 0.6% glucose, 2 mM GlutaMAX (Thermo Fisher Scientific), 2% B27 Plus (Thermo Fisher Scientific), and 100 U/mL Penicillin-Streptomycin (Thermo Fisher Scientific). These cells were kept at 37°C in a humidified atmosphere of 95% air and 5% CO_2_.

### Immunocytochemistry

Cells were washed in PBS at 37°C and then fixed in 4% paraformaldehyde for 20 minutes. Subsequently, cells were permeabilized with 0.1% Triton X-100 for 3 minutes and then blocked with 5 or 10% bovine serum albumin (BSA, Merck) in PBS for 20 minutes. Antibodies were diluted in Can Get Signal Solution (Toyobo). Proteins were probed with primary antibodies for 1 hour at room temperature. Subsequently, they were incubated with secondary antibodies for 1 hour at room temperature. Actin visualized with Alexa Fluor 555 Phalloidin (Thermo Fisher Scientific, 1:200) at the same time as probing with secondary antibody.

For detection of antigens on the membrane surface of live cells, anti-HA tag antibody was diluted at 1:250 in a culture medium and reacted for 30 minutes in a cell culture incubator. In addition, PFA fixation, detergent permeabilization, blocking treatment, other primary antibody reactions were carried out as usual, and secondary antibody was reacted.

Using the following procedure, the molecules that do not interact with the cytoskeleton were removed. First, a sufficient amount of saponin (Kanto Chemical) was dissolved in water to make a saturated solution. Then, the saturated solution of saponin was added to the culture medium to reach a final concentration of 0.1% and permeabilized in the incubator for 3 minutes. After permeabilization, the cells were immediately immersed in methanol at -30°C and fixed on ice. After that, additional permeabilization with Triton X-100 was performed. The rest of the procedure was the same as for normal immunofluorescence staining.

Immunofluorescence staining images were mainly acquired with a confocal laser scanning microscope (LSM 880; ZEISS) equipped with a 63x/1.4 Plan Apochromat oil immersion objective. For statistical analysis of COS-7 morphology, cells were observed with an IX71 inverted microscope (Olympus) equipped with DeltaVision (Cytiva), CoolSNAP HQ2 CCD camera (Teledyne Photometrics), and 40x/1.35 UApo/340 oil immersion objective. Tile imaging was performed by randomly selecting a region that contained approximately 100-300 EGFP-expressing cells, and cells with more than 10 protrusion-like structures with branches larger than 10 μm were manually counted for statistical analysis. Detailed observation of the cytoskeleton was performed on an Eclipse Ti2 inverted microscope (Nikon) equipped with a Dragonfly spinning disk (Oxford Instruments), an EMCCD iXon Ultra 897 (Oxford Instruments), and a 60x/1.4 Plan Apo oil immersion objective. After image acquisition, super-resolution images were created by SRRF-Stream.

### Antibodies

An affinity-purified rabbit polyclonal antibody specific for TEN2 was raised against a synthetic peptide of the sequence CSNTSHQIMDTNPDE (Eurofins Genomics). This is a cysteine added to the C-terminus of amino acids 203-216 in the intracellular domain. After crude purification of serum with ammonium sulfate, it was purified by affinity chromatography using SulfoLink Coupling Resin (Thermo Fisher Scientific) coupled with a synthetic peptide of the sequence CQMPLLDSNTSHQIMDTNPDEEFSPNS (GenScript). This is amino acids 196-222 in the intracellular domain. For other antibodies, commercially available antibodies were used as shown in the table (Table S7).

### Quantification of gephyrin puncta

Gephyrin was visualized using anti-gephyrin. For dendritic tracing, dendrites were visualized by expressing TagRFP in knockdown experiments and by immunofluorescence staining with MAP2 in other experiments. TagRFP or MAP2 channels were binarized to clarify dendrites and traced using the NeuronJ plugin. The number of gephyrin puncta was quantified by SynapCountJ plugin using this trace data and gephyrin channels. Since this plugin requires two images, we made another copy of the gephyrin channel and identified the gephyrin puncta by the same threshold.

### Cluster analysis and statistical analysis

Cluster analysis and statistical analysis were performed using Excel (Microsoft) and R software. Statistical tests, sample sizes, and experiments are shown in the figure legends. In the cluster analysis, the number of clusters was set to 3 after a hierarchical cluster analysis was conducted without pre-determining the number of clusters in the preliminary experiment. In the main experiment, the number of clusters was set to 3 in advance and then analyzed. The cluster analysis did not use statistical methods to determine sample size a priori, but instead targeted all postsynapses within 100 μm of the cell body of the eight neurons. Postsynapses were detected at a threshold of 32768 with 16 bits of gephyrin fluorescence intensity taken with the same criteria. The ratio of neurons with biased MAP2 localization was preplanned to obtain statistics from three independent experiments. For the other experiments, the number of samples was estimated with the “power.t.test” and “power.anova.test” functions in R so that power = 0.8 from the first experiment (n = 4 ∼ 6), and observations of the set number of samples were made in three or more independent biological samples. Even when the set number was exceeded, samples were not excluded and all observed samples were used for statistics. The number of samples was not re-set from the final power. Before the significance tests, the Kolmogorov-Smirnov test was used to test for normal distribution. Nonparametric methods were used to analyze samples that were not normally distributed. t-tests were based on the Welch method and did not assume equal variance. The experimenter was not blinded to the experimental conditions during all data acquisition or quantification.

### Motif search and alignment

The columns containing the Uniprot IDs of proteins from the table of Excitatory and Inhibitory Synaptic Cleft Proteomes (Loh et al., 2016) were converted to comma-separated csv files and then converted to fasta format using NCBI E-utilities. Fuzzpro in the EMBOSS package (Rice et al., 2000) were used for each motif search for the following conditions: pattern Sx[ILV]P, mismatch 0 for SxφP motif; pattern LRPPTP[ILV], mismatch 2 for LxxPTPφ motif. This search was performed in a macOS terminal with Anaconda installed. The obtained sequences were searched with Uniprot and manually checked whether they were extracellular or cytosolic. The alignment of amino acid sequences was performed using Clustal Omega (EMBL-EBI) and MacVector (MacVector).

### Plasmids

For the full-length EGFP-Teneurin2 clone, partial Teneurin2 from KAZUSA cDNA (NCBI AB032953) and RIKEN cDNA (NCBI AK031198) were amplified by PCR and inserted into pEGFP-C1 (Takara Bio). The missing part was complemented by a custom gene synthesis (Eurofins Genomics K.K.). The protein translated from this plasmid is equivalent to the full-length Homo sapiens Teneurin-2 (NCBI NP_001382389), consisting of 2774 amino acids. Note that there are six mutations in our construction as follows: I418V, M431V, V590L, S659A, T720S and L2384P. They are not located in the intracellular domain. EB1-TagRFP was generated by PCR amplification from KAZUSA cDNA (NCBI AB463888) and inserted into pTagRFP-N (Evrogen). The shRNA target sequence was designed for protein knockdown using the BLOCK-iT RNAi Designer tool (Thermo Fisher Scientific). A cassette containing the pre-shRNA sequence was inserted into pBAsi-mU6 (Takara Bio). The target sequences of each shRNA are as follows: Negative control, GCCTAAGGTTAAGTCGCC; Teneurin2 #1, GCCAGGTTTGATTATACC. For volume marker, the SV40 promoter and tagRFP sequences were amplified and inserted into the pBAsi-mU6 vector.

### Transfection

COS-7 cells were transfected with the plasmid using Lipofectamine 2000 (Thermo Fisher Scientific). Transfection conditions are described in each experiment. According to the manufacturer’s protocol, the cultured neurons were transfected using the High-Efficiency Ca^2+^ Phosphate Transfection Kit (Takara Bio). Briefly, the culture medium was replaced with fresh MEM containing pyruvate, glucose, and GlutaMAX. 2 µg of plasmid, 3.1 µl of 2 M CaCl_2_, and 25 µl of Hanks equilibrium salt solution were mixed by pipetting and vortexing and incubated at room temperature for 15 minutes. Next, the DNA/Ca^2+^ phosphate suspension was added to the culture medium and incubated in a 5% CO_2_ incubator at 37°C for 1 hour. After the incubation, the DNA/Ca^2+^ phosphate precipitates were dissolved for 15 minutes with the pre-equilibrated medium in a 10% CO_2_ incubator and then replaced with the original medium.

### Quantitative analysis of correlation coefficient

For the analysis of TEN2-EB1 colocalization, COS-7 cells were plated in a Lab-Tek II 8-well chamber coverglasses at 1-2 x 10^5^ /well. 0.5 μg of pEGFP-C1 vector, which inserted the necessary part of TEN2, and 1 μg of EB1-TagRFP, and 0.5 μL Lipofectamin 2000 were diluted in Opti-MEM (Thermo Fisher Scientific) and transfected for 1 well. After 18 hours, cells were fixed with PFA and cells expressing EB1 in MTs pattern were recorded with LSM 880. To exclude regions with no signal and full of noise, EGFP channels were binarized to detect cell morphology, and these were used as regions of interest (ROI). Since the background was sufficiently reduced under these conditions, quantitative analysis was performed using the non-threshold Pearson’s R value as the correlation coefficient between TEN2 and EB1.

For the analysis of TEN2-MAP2 colocalization, neurons expressing EGFP-TEN2 were immunostained with MAP2. The signal intensities of EGFP and MAP2 perpendicular to the direction of dendrite elongation were quantified using ImageJ’s Plot Profile, and the correlation coefficient between the two variables was calculated using Excel’s CORREL function for quantitative analysis.

### Coating of anti-HA tag antibody

Lab-Tek II 8-well chamber coverglasses were coated with poly-D-lysine overnight and washed twice with sterile water. The additional coating was done overnight with HBSS containing rabbit monoclonal anti-HA tag (Cell Signaling Technology, #3724) diluted 1:20. After washing twice with sterile water, the chamber was immediately plated with COS-7 with DMEM with no attempt to dry the chamber. Transfection was performed within 8 hours.

### TIRF live imaging and analysis of EB1 comet

COS-7 cells were plated in a Lab-Tek II 8-well chamber coverglasses at 1-2 x 10^5^ /well. 0.5 μg of pEGFP-C1 vector, which inserted the necessary part of TEN2, and 0.05 μg of EB1-TagRFP, and 0.5 μL Lipofectamin 2000 were diluted in Opti-MEM and transfected for 1 well. One hour after transfection, the medium was changed to fresh medium, and observations were carried out 12-18 hours later. At the time of observation, the medium was replaced with Leibovitz L-15 medium (Wako). TIRF observation was performed using an Eclipse Ti inverted microscope (Nikon) equipped with a 488-nm and 568-nm sapphire laser (Coherent), a Plan Apo TIRF 100x Oil Immersion objective, and a house-made thermostatic chamber to maintain temperature at 37°C during observation. For time-lapse recording, images were acquired using an EMCCD iXon3 DU897 (Oxford Instruments) with an exposure time of 1.0 second and a gain of 100, taking 50 images every 2 seconds for 1 minute 38 seconds. The perfect focus system (PFS) was used to maintain focus during recording.

The recorded images were analyzed using Particle Tracker 2D/3D, one of the Mosaic plugins in ImageJ, to detect trajectories under the following conditions: radius = 3, cutoff = 3, per/abs = 0.4, link = 1, displacement = 5, dynamics = straight. Each trajectory’s velocity was calculated by dividing the distance moved per frame by the interval time of 2 seconds, and the average of the velocities over the number of frames of the trajectory was defined as the velocity of the corresponding EB1 comet.

### CRISPR/Cas9-mediated knock-in of the 3xHA tag into Tenm2 gene

The Tenm2 3xHA tag knock-in mice were generated by using a CRISPR/Cas9 genome-editing technology onto pronuclear stage embryos (Doudna & Charpentier, 2014). In brief, female C57BL/6J JAX mice (Charles River; IMSR Cat# JAX:000664, RRID:IMSR_JAX:000664) were super-ovulated by intraperitoneal injection of 7.5 units of pregnant mare’s serum gonadotropin (PMSG; ASKA Pharmaceutical), followed by 7.5 units of human chorionic gonadotropin (hCG; ASKA Pharmaceutical) 48 hours later. Fifteen hours after the hCG injection, super-ovulated female mice were euthanized via cervical dislocation, and unfertilized oocytes isolated from the female mice were subjected to in vitro fertilization with freshly isolated spermatozoa from euthanized C57BL/6J JAX male mice, as previously described (Kaneko et al., 2018). Introduction of Cas9 protein, guide RNA, and single strand oligodeoxynucleotide (ssODN) into pronuclear stage embryos was carried out using the TAKE method (Kaneko, 2017). Cas9 protein, guide RNA, and ssODN were purchased from IDT (Integrated DNA Technologies). Mixture of crRNA and tracrRNA was used as guide RNA. Guide RNA and ssODN were designed to insert 3xHA tag sequences just upstream from the stop codon of the Tenm2 gene of the C57BL/6 mouse (guide RNA: 5’-GACAGAATGAGATGGGAAAG-3’, ssODN: 5’-ACAGTAGCAGCAACATCCAGTTCTTAAGACAGAATGAGATGGGAAAGAGATACCCATACGATGTACCTGACTATGCGGGCTATCCCTATGACGTCCCGGACTATGCAGGATCCTATCCTTATGACGTTCCAGATTACGCTGTTTAACAAAATAACCTGCTGCCACCTCTTCTCTGGGTGGCTCAGCAGGAGCAACT-3’, where 3xHA tag sequences are underlined). The CRISPR/Cas9 solution contained 50 ng/μL Cas9 protein, 50 ng/μL crRNA, 50 ng/μL tracrRNA, and 100 ng/μL ssODN in Opti-MEM (Thermo Fisher Scientific). Super electroporator NEPA21 (NEPA GENE) was used to introduce Cas9 protein, guide RNA, and ssODN into embryos. The poring pulse was set to voltage: 225 V, pulse length: 2.0 ms, pulse interval: 50 ms, number of pulses: 4, decay rate: 10%, polarity: +. The transfer pulse was set to a voltage: 20 V, pulse length: 50 ms, pulse interval: 50 ms, number of pulse: 5, decay rate: 40%, Polarity: +/-. The CRISPR/Cas9 solution (45 μL) was filled between metal plates of 5 mm gap electrodes on a glass slide (CUY505P5, NEPA GENE). The embryos placed in line between the electrodes were then discharged. The embryos were then cultured in HTF at 37 °C in 5% CO_2_/95% air. On the next day, two-cell embryos were transferred into the oviduct ampulla (40–48 embryos per oviduct) of pseudopregnant ICR (Japan SLC; MGI Cat# 5462094, RRID:MGI:5462094) females. All mice generated were genotyped by PCR amplification of genomic DNA isolated from the tip of tail, followed by sequencing. Sequence of the primers used for genotyping were as follows; Tenm2ex29+564F; CAAGGAGCAGCAGAAAGCCAG; Tenm2ex29+871R; TAAAGCAGCCCGGCCTCAGTG. The resulting PCR product was cut by BamHI, and the expected size was 308 bp for wild-type and 254 bp and 147 bp for Tenm2 3xHA tag knock-in mice.

Mice were backcrossed with wild-type C57BL/6J at least four times, and at least one of them was with a wild-type male to replace the Y chromosome. Mice were kept in an environment free of specific pathogens according to the institutional guidelines of Gunma University. All mice have been genotyped by PCR. These experiments have passed a rigorous ethical review and have been approved by Gunma University for animal experiments (approval number: 20-061) and genetic recombination experiments (approval number: 21-042).

### Proximity ligation assay

The ploximity ligation assay uses DuoLink (Merck) and follows the protocol distributed by the supplier. Primary antibodies were reacted according to the immunocytochemistry protocol described above. PLA Probe Anti-Mouse PLUS (Merck) and PLA Probe Anti-Rabbit MINUS (Merck) diluted 1:10 and reacted at 37°C for 1 hour. After three washes, oligonucleotides labeled with the two secondary antibodies were ligated with ligase, DNA was amplified by rolling circle amplification, and the Duolink in situ detection reagent green (Merck) was incorporated into the newly synthetic DNA. After three washes and a 5-minute post-fix, the neurons were reacted with a secondary antibody against MAP2 for 1 hour. Observations were made after three additional washes.

### dSTROM

The equipment used in dSTORM was the same setup as the Nikon-Ti microscope used in the TIRF live imaging described above. Immunofluorescence staining for dSTORM was performed using a concentration of both primary and secondary antibodies that was four times higher than that of normal immunofluorescence staining, but other details were the same as for normal immunofluorescence staining. Observations were made in 50 mM Tris-HCl (pH8.0), 10 mM NaCl buffer containing 0.1 M MEA, 0.7 mg/ml glucose oxidase, 10% glucose and 0.034 mg/ml catalase. Observations were taken continuously at 59 Hz for AF647, CF568, and AF488, in that order, and 25000 images each were recorded for the first two dyes; AF488 does not blink under these conditions, so only 500 images were recorded and used only to determine if AF488 was positive or negative. All recordings were drift-corrected and then STORM images were constructed using Nikon’s accompanying analysis software, NIS.

## Author contributions

S.I. and H.I. conceived the study. S.I. and Y.S. performed the cell biological experiments. R.K. generated knock-in mice. S.I. and H.I. analyzed the data and discussed and wrote the paper.

## Acknowledgements

We thank Drs. Tohru Murakami and Yuki Tajika. We thank Yoshihiro Morimura, Toshie Kakinuma, and Sachiko Sato for technical assistance and care of the mice. This work was supported by a Grant-in-Aid for Scientific Research (C) (18K06499, 22K06805, H.I) and Young Scientist (20K16104, S.I.) from the Ministry of Education, Culture, Sports, Science and Technology of Japan, and Takeda Science Foundation. This work used equipment shared in the MEXT Project for promoting public utilization of advanced research infrastructure (JPMXS0420600119, JPMXS0420600120, JPMXS0420600121).

## Materials Availability

Further information and requests for resources and reagents should be directed to and will be fulfilled by the lead contact, Hirohide Iwasaki.

## Video Legends

**Figure 3–video 1. TEN2TM does not capture dynamic MTs**

Time-lapse observation of EB1 comet in COS-7 cells expressing TEN2TM without immobilization as control.

**Figure 3–video 2. Immobilized TEN2 captures dynamic MTs**

Time-lapse observation of EB1 comet in COS-7 cells expressing TEN2N-L immobilized by antibodies. Slower or almost immobile EB1 comets are shown with arrows.

## Source Data Legends

**Figure 1–source data 1**

4 Excel sheets containing the numerical data used to generate the figures 1E, H, I and J.

**Figure 1–figure supplement 1–source data 1**

2 Excel sheets containing the numerical data used to generate the figures 1–figure supplement 1B, C and F. The data used for B and C are combined in one file.

**Figure 1–figure supplement 2–source data 1**

2 Excel sheets containing the numerical data used to generate the figures 1–figure supplement 2C and E.

**Figure 2–source data 1**

2 Excel sheets containing the numerical data used to generate the figures 2D and F.

**Figure 3–source data 1**

3 Excel sheets containing the numerical data used to generate the figures 3E, G and H.

**Figure 4–source data 1**

3 Excel sheets containing the numerical data used to generate the figures 4H, I and J.

**Figure 4–figure supplement 1–source data 1**

Unprocessed full-size gel photograph showing genotyping of TEN2 knock-in mice and photograph showing the region used in figures 4–figure supplement 1C with dashed lines.

**Figure 4–figure supplement 1–source data 2**

An Excel sheet containing the numerical data used to generate the figure 4–figure supplement 1E.

**Figure 5–source data 1**

2 Excel sheets containing the numerical data used to generate the figures 5B and D.

**Figure 5–figure supplement 1–source data 1**

An Excel sheet containing the numerical data used to generate the figure 5–figure supplement 1D.

**Figure 6–source data 1**

4 Excel sheets containing the numerical data used to generate the figures 6B, C, D and F.

**Figure 1–figure supplement 1.**
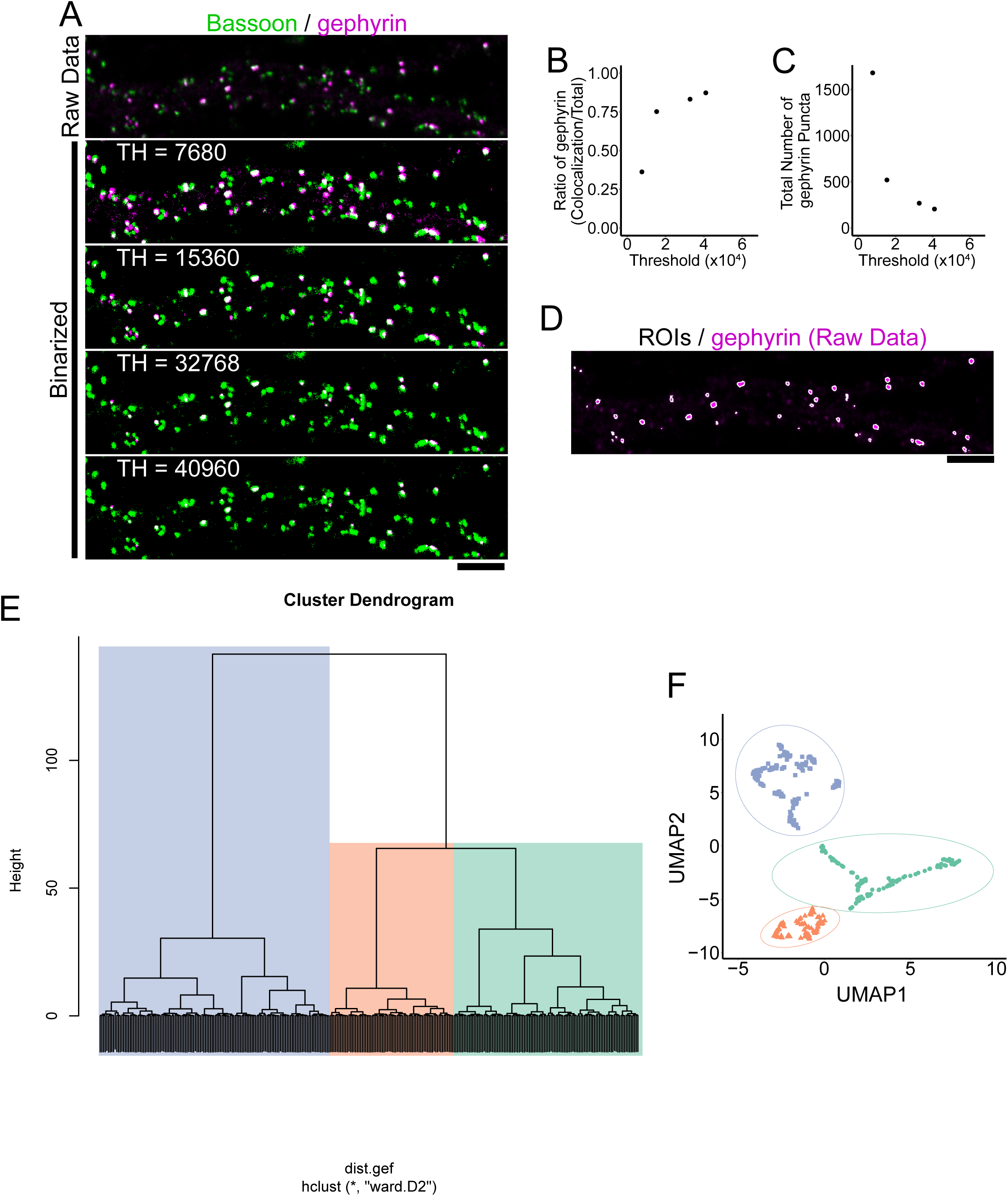
Cluster analysis of inhibitory postsynapses and correlation with TEN2. (A) Image of immunofluorescence staining of bassoon and gephyrin in DIV20 hippocampal cultured neuron. Bassoon is binarized with a threshold value of 32768. Varying the threshold for gephyrin binarization changes the ratio of colocalization. Scale bar, 5 μm. (B) Plots showing the threshold for gephyrin binarization and the colocalization ratio with bassoon. Lowering the threshold lowers the colocalization ratio because more gephyrin is detected. When the threshold is high, more than 80% of gephyrin colocalizes with bassoon. (C) Plot of gephyrin binarization threshold and number of gephyrin punctures detected. At lower thresholds, more gephyrins are detected. Increasing the threshold decreases the number of gephyrins detected, but the slope of the decrease is slower. (D) ROIs (= postsynaptic regions) obtained by binarization with gephyrin threshold set to 32768 and superimposition of raw data. Human-recognizable postsynapses are detected almost as intuitively. (E) Preliminary experiments for setting the number of clusters. Three parameters, synaptic area, MAP2 intensity, and actin intensity, were reduced to two dimensions by UMAP and then analyzed hierarchically by Ward’s method. Based on the results, it was determined that it was appropriate to divide the data into three clusters. (F) Results of cluster analysis in preliminary experiments. The number of synapses belonging to each cluster was 111, 138 and 74, observed by three independent experiments.

**Figure 1–figure supplement 2.**
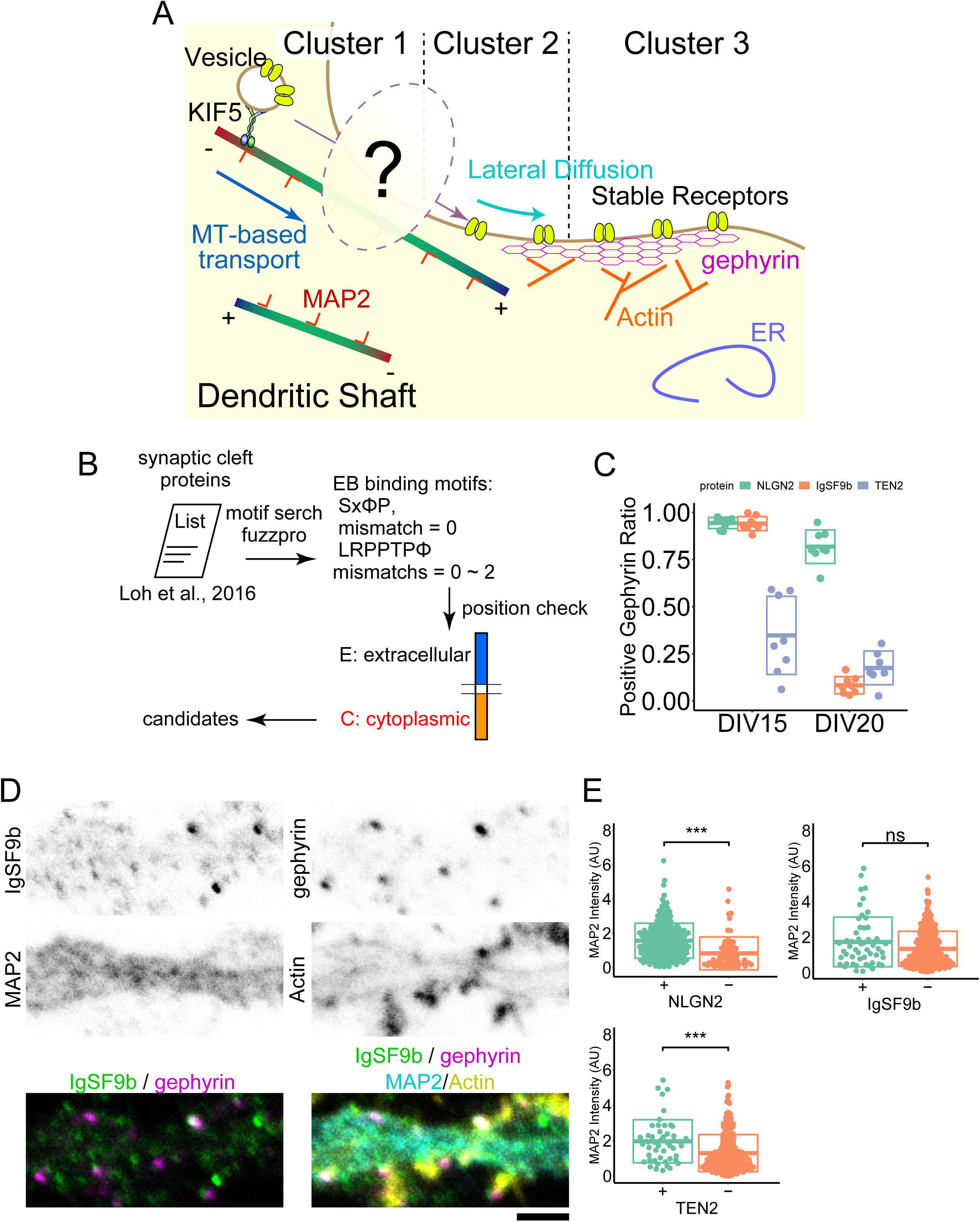
Cluster analysis of inhibitory postsynapses and correlation with TEN2. (A) Assumptions of the situation reflected by clustering. Cluster 3 belongs to a stable postsynapse anchored by gephyrin and actin. Cluster 1 belongs to a dynamic postsynapse with receptors being moved in and out by an MTs-based transport system. Cluster 2 belongs to an intermediate in which intense lateral diffusion is taking place. How the transition between clusters 1 and 2 occurs has been unclear to date. (B) Schematic diagram of motif search. Two types of EB1 binding motifs were searched from the list of molecules obtained from the proteome analysis. Whether the searched motifs were extracellular or cytoplasmic was manually checked. (C) Ratio of gephyrin puncta positive for each adhesion molecule. The ratio of positive for each molecule decreased in DIV20 compared to DIV15, but the decrease was particularly prominent for IgSF9b. mean ± SD at DIV15: NLGN2, 0.94 ± 0.03; IgSF9b, 0.94 ± 0.04; TEN2, 0.34 ± 0.21. mean ± SD at DIV20: NLGN2, 0.82 ± 0.09; IgSF9b, 0.08 ± 0.05; TEN2, 0.17 ± 0.09. The number of observations was 8, 8, 8, 8, 8 and 7 neurons, respectively. (D) Images of immunofluorescence staining of IgSF9b, gephyrin, MAP2, and actin in DIV20 hippocampal cultured neuron. Scale bar, 2 μm. (E) Classical comparative quantification of MAP2 intensity without reflecting cluster analysis. Welch’s t-test results showed a significant difference between positive and negative synapses for NLGN2 (p = 4.08e^-07^) and TEN2 (p = 6.48e^-04^), but not for IgSF9b (p = 0.059). The number of NLGN2 positive and negative synapses are 228 and 65. The number of IGSF9b positive and negative synapses are 53 and 283. The number of TEN2 positive and negative synapses are 49 and 262. ***p < 0.001. ns, not significant.

**Figure 4–figure supplement 1.**
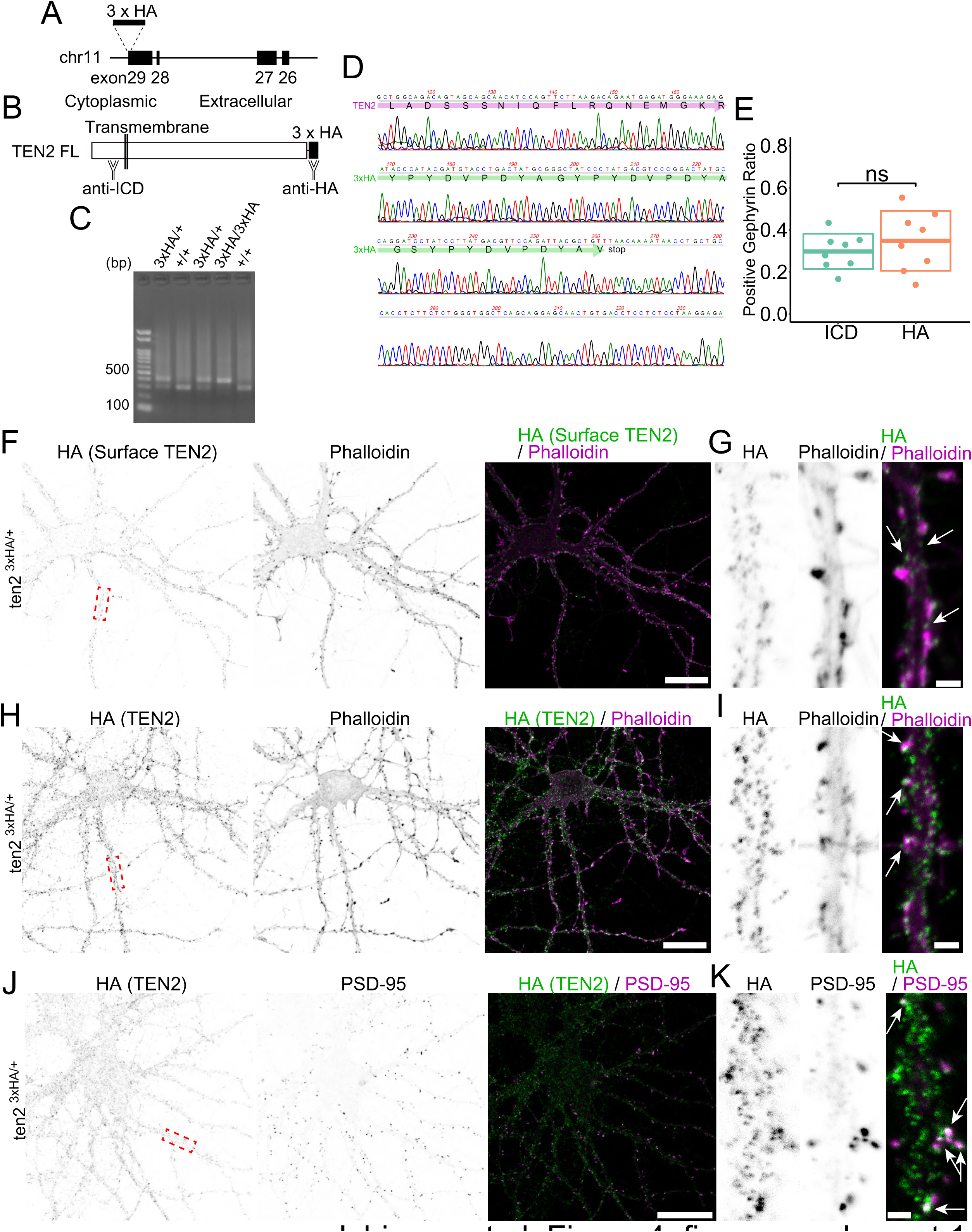
TEN2 localization at the semi-periphery region of the inhibitory postsynapse. (A) Overview of knock-in mice. TEN2 is encoded on the minus strand of chromosome 11. Knock-in mice were generated by inserting a 3xHA sequence before the stop codon in Exon29. (B) Overview of the teneurin-2 full-length protein and antibody recognition sites. Teneurin is a type II transmembrane protein that is intracellular at its N-terminus and extracellular at its C-terminus. 3xHA, inserted just before the stop codon, is extracellular when translated. (C) Typical genotyping results. A 300 bp band is seen in wild-type mice, while a +100 bp band is seen in knock-in mice; if both bands are seen, the mouse is heterozygous with only one allele being knock-in. (D) Sequence confirmation by Sanger sequencing. Bands amplified by genotyping were purified and Sanger sequenced to confirm the knock-in sequence. (E) No effect of HA knock-in on localization to inhibitory synapses. ICR-delivered wild-type neurons at DIV15 were co-stained with ICD and gephyrin antibodies, and HA knock-in neurons at DIV15 were co-stained with HA and gephyrin antibodies. mean ± SD were 0.30 ± 0.01 and 0.34 ± 0.02, respectively. Since there was no significant difference in the ratio of colocalization (p = 0.40), we concluded that HA knock-in had no effect on localization to inhibitory synapses. (F) Images of immunofluorescence staining of HA tag and actin exposed on the cell membrane surface in the knock-in neuron. The red dashed box is magnified in (G). (G) Confirmation that HA tag is exposed at the plasma membrane surface, suggesting that TEN2 functions at the plasma membrane surface. (H) Images of immunofluorescence staining of HA tag and actin in the knock-in neuron. The red dashed box is magnified in (I). (I) HA tag signals are also present in the dendritic shaft but are particularly strong near spine-like structures, suggesting that the molecule is more abundant at excitatory synapses. Arrows, representative colocalization. (J) Images of immunofluorescence staining of HA tag and PSD-95 in the knock-in neuron. The red dashed box is magnified in (K). (K) The strong signal of HA tag at the site where PSD-95 is localized suggests that the molecule is abundant at excitatory synapses. Arrows, representative colocalization. Scale bars indicate 50 μm in (F), (H), and (J) and 2 μm in (G), (I), and (K).

**Figure 5–figure supplement 1.**
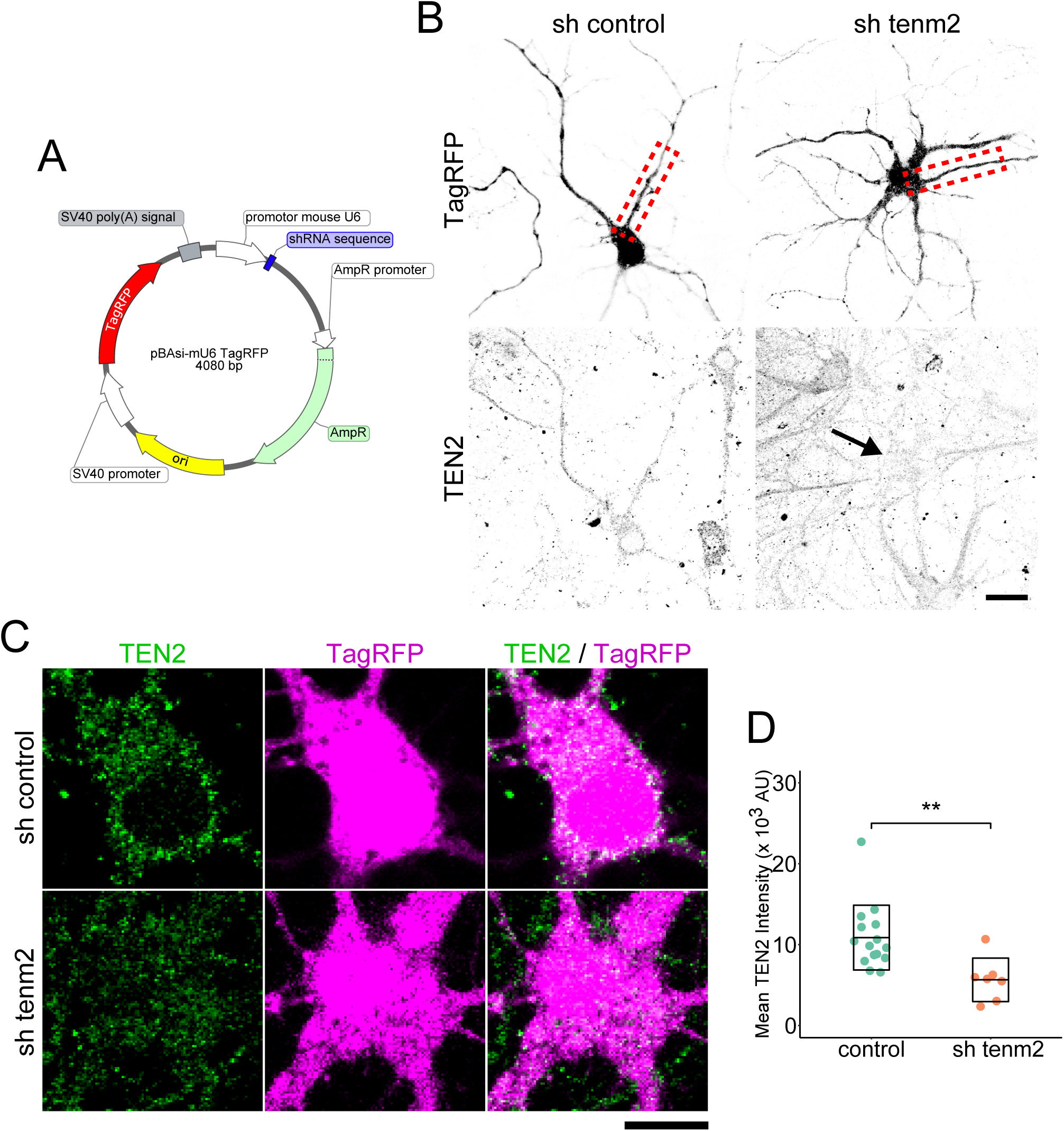
Inhibitory postsynapse maturation induced by postsynaptic TEN2. (A) Overview of the knockdown vector. The shRNA sequence and TagRFP expression gene are located downstream of the mouse U6 promoter and SV40 promoter, respectively. (B) Low-magnification images of knocked-down neurons and confirmation of TEN2 knockdown. The knockdown vector-transfected neurons were stained for endogenous TEN2 and gephyrin. The TEN2 signal was reduced in the TEN2 knockdown neurons compared to the surrounding or control neurons (arrows). Red dashed boxes are shown in Figure. 5A. The scale bar indicates 20 μm. (C) High magnification images of knocked-down neurons. Neurons transfected with knockdown vector were stained with endogenous TEN2. A reduction of TEN2 signal is observed in knockdown neurons. Scale bar indicates 10 μm. (D) Plots and crossbars (mean ± SD) showing quantification of TEN2 signal intensity in cells transfected with knockdown vectors. The mean signal intensity of TEN2 was 10.9 ± 4.00 x 10^3^ arbitrary units (AU) in neurons transfected with the control vector, and 5.65 ± 2.70 x 10^3^ AU in neurons transfected with TEN2 knockdown vector (p = 0.0022). n = 15 control neurons and 7 knockdown neurons, from three independent experiments. **p < 0.01.

**Table S1.**
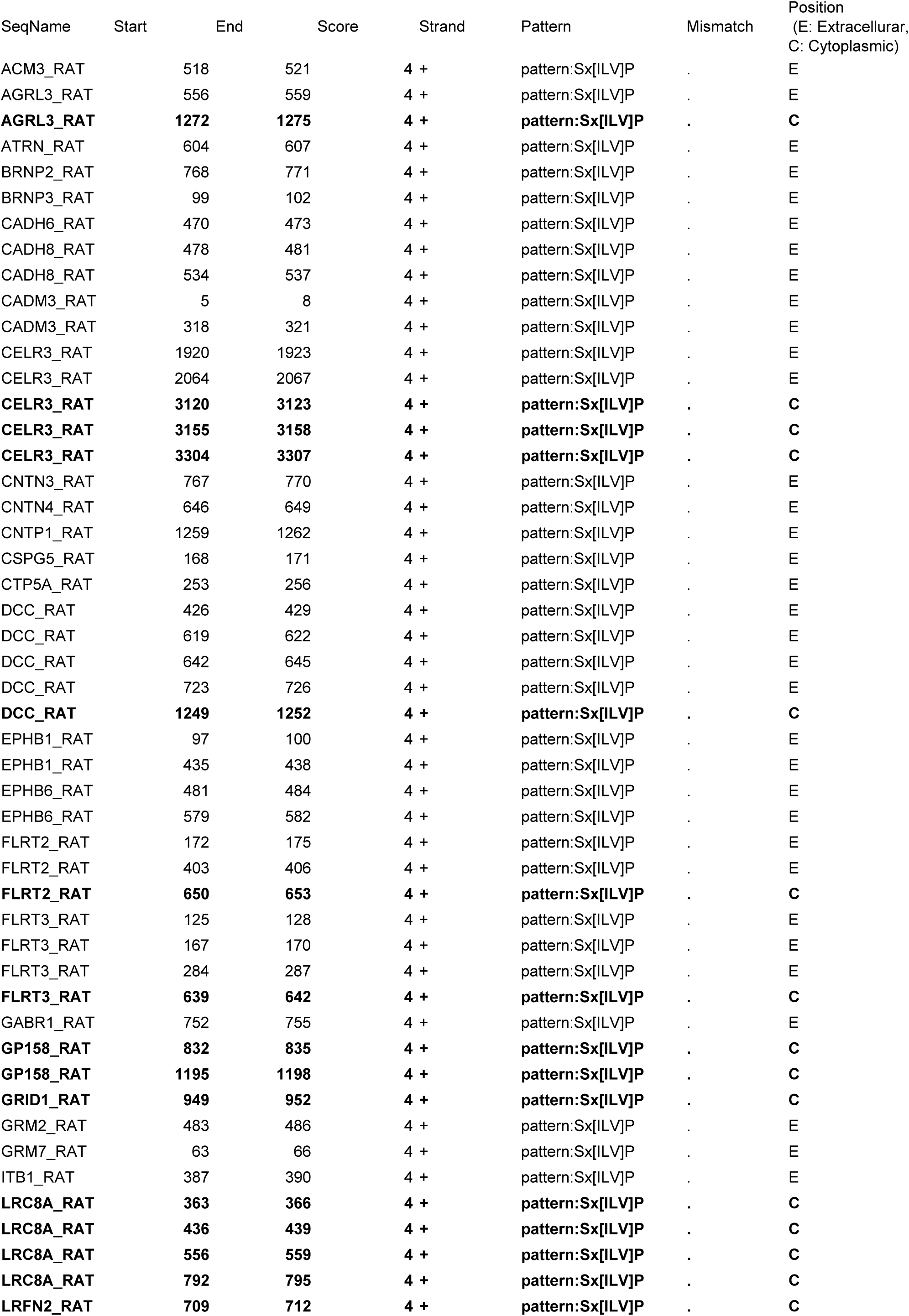
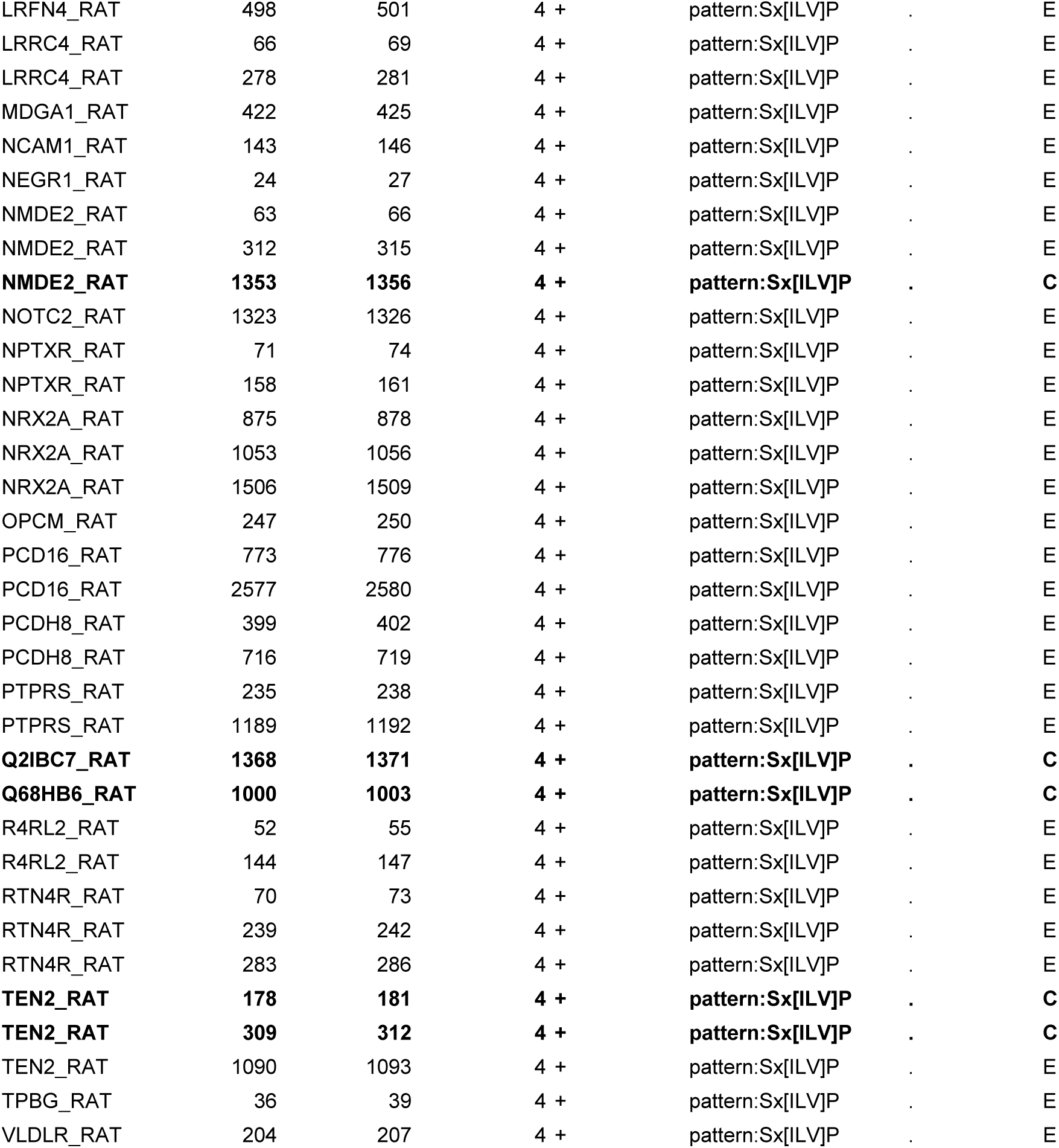
The excitatory synaptic cleft proteins with SxφP motif related to Figure 1

**Table S2.**
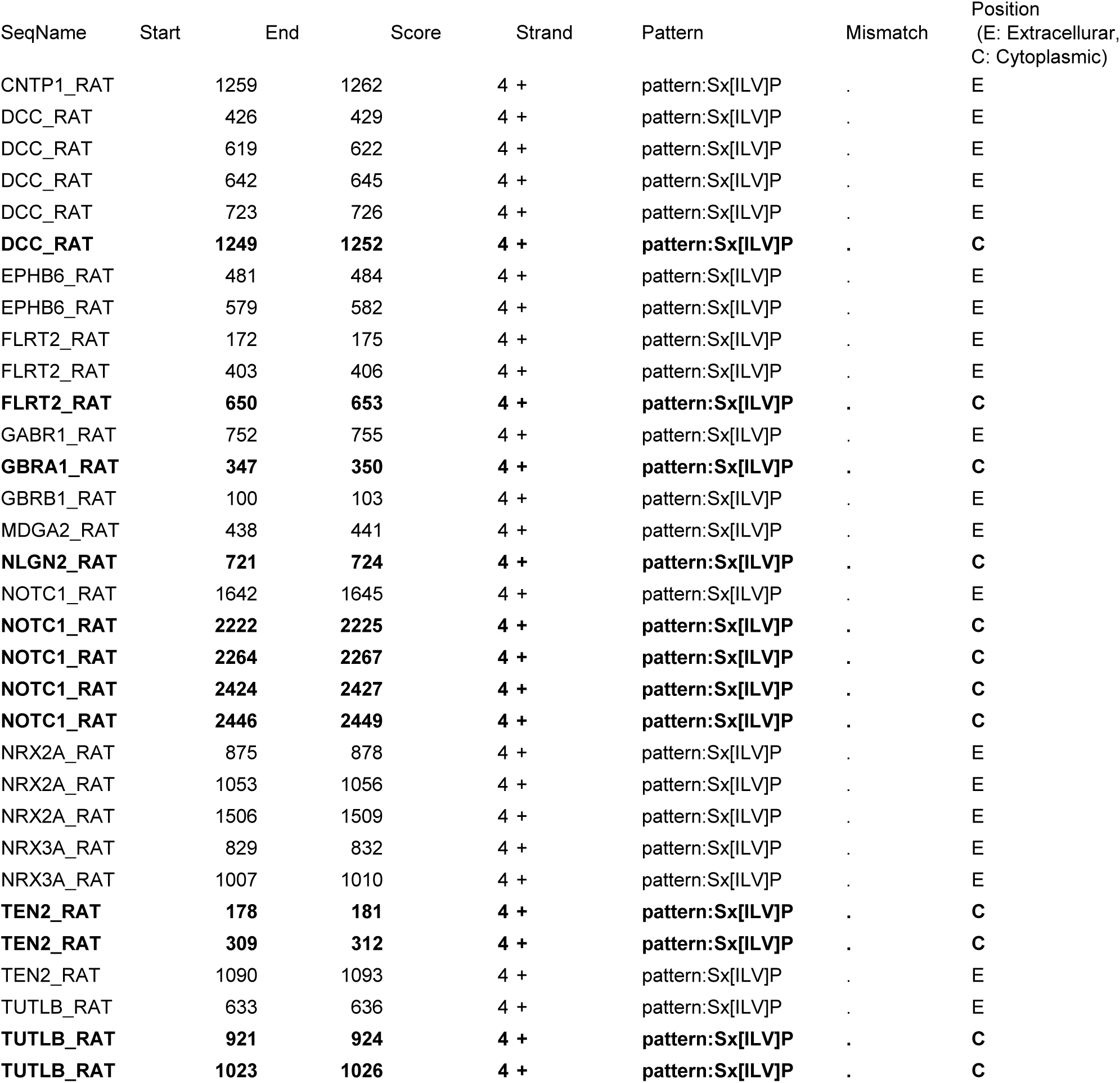
The inhibitory synaptic cleft proteins with SxφP motif related to Figure 1

| SeqName | Start | End | Score | Strand | Pattern | Mismatch | Position<br>(E: Extracellular,<br>C: Cytoplasmic) |
| --- | --- | --- | --- | --- | --- | --- | --- |
| CNTP1_RAT | 1259 | 1262 | 4 + |  | pattern:Sx[ILV]P | . | E |
| DCC_RAT | 426 | 429 | 4 + |  | pattern:Sx[ILV]P | . | E |
| DCC_RAT | 619 | 622 | 4 + |  | pattern:Sx[ILV]P | . | E |
| DCC_RAT | 642 | 645 | 4 + |  | pattern:Sx[ILV]P | . | E |
| DCC_RAT | 723 | 726 | 4 + |  | pattern:Sx[ILV]P | . | E |
| **DCC_RAT** | **1249** | **1252** | **4 +** |  | **pattern:Sx[ILV]P** | . | **C** |
| EPHB6_RAT | 481 | 484 | 4 + |  | pattern:Sx[ILV]P | . | E |
| EPHB6_RAT | 579 | 582 | 4 + |  | pattern:Sx[ILV]P | . | E |
| FLRT2_RAT | 172 | 175 | 4 + |  | pattern:Sx[ILV]P | . | E |
| FLRT2_RAT | 403 | 406 | 4 + |  | pattern:Sx[ILV]P | . | E |
| **FLRT2_RAT** | **650** | **653** | **4 +** |  | **pattern:Sx[ILV]P** | . | **C** |
| GABR1_RAT | 752 | 755 | 4 + |  | pattern:Sx[ILV]P | . | E |
| **GBRA1_RAT** | **347** | **350** | **4 +** |  | **pattern:Sx[ILV]P** | . | **C** |
| GBRB1_RAT | 100 | 103 | 4 + |  | pattern:Sx[ILV]P | . | E |
| MDGA2_RAT | 438 | 441 | 4 + |  | pattern:Sx[ILV]P | . | E |
| **NLGN2_RAT** | **721** | **724** | **4 +** |  | **pattern:Sx[ILV]P** | . | **C** |
| NOTC1_RAT | 1642 | 1645 | 4 + |  | pattern:Sx[ILV]P | . | E |
| **NOTC1_RAT** | **2222** | **2225** | **4 +** |  | **pattern:Sx[ILV]P** | . | **C** |
| **NOTC1_RAT** | **2264** | **2267** | **4 +** |  | **pattern:Sx[ILV]P** | . | **C** |
| **NOTC1_RAT** | **2424** | **2427** | **4 +** |  | **pattern:Sx[ILV]P** | . | **C** |
| **NOTC1_RAT** | **2446** | **2449** | **4 +** |  | **pattern:Sx[ILV]P** | . | **C** |
| NRX2A_RAT | 875 | 878 | 4 + |  | pattern:Sx[ILV]P | . | E |
| NRX2A_RAT | 1053 | 1056 | 4 + |  | pattern:Sx[ILV]P | . | E |
| NRX2A_RAT | 1506 | 1509 | 4 + |  | pattern:Sx[ILV]P | . | E |
| NRX3A_RAT | 829 | 832 | 4 + |  | pattern:Sx[ILV]P | . | E |
| NRX3A_RAT | 1007 | 1010 | 4 + |  | pattern:Sx[ILV]P | . | E |
| **TEN2_RAT** | **178** | **181** | **4 +** |  | **pattern:Sx[ILV]P** | . | **C** |
| **TEN2_RAT** | **309** | **312** | **4 +** |  | **pattern:Sx[ILV]P** | . | **C** |
| TEN2_RAT | 1090 | 1093 | 4 + |  | pattern:Sx[ILV]P | . | E |
| TUTLB_RAT | 633 | 636 | 4 + |  | pattern:Sx[ILV]P | . | E |
| **TUTLB_RAT** | **921** | **924** | **4 +** |  | **pattern:Sx[ILV]P** | . | **C** |
| **TUTLB_RAT** | **1023** | **1026** | **4 +** |  | **pattern:Sx[ILV]P** | . | **C** |

**Table S3.** The excitatory and inhibitory common synaptic cleft proteins with SxφP motif related to Figure 1

| SeqName | Start | End | Score | Strand | Pattern | Mismatch | Position<br>(E: Extracellular,<br>C: Cytoplasmic) |
| --- | --- | --- | --- | --- | --- | --- | --- |
| CNTP1_RAT | 1259 | 1262 | 4 + |  | pattern:Sx[ILV]P | . | E |
| DCC_RAT | 426 | 429 | 4 + |  | pattern:Sx[ILV]P | . | E |
| DCC_RAT | 619 | 622 | 4 + |  | pattern:Sx[ILV]P | . | E |
| DCC_RAT | 642 | 645 | 4 + |  | pattern:Sx[ILV]P | . | E |
| DCC_RAT | 723 | 726 | 4 + |  | pattern:Sx[ILV]P | . | E |
| **DCC_RAT** | **1249** | **1252** | **4 +** |  | **pattern:Sx[ILV]P** | . | **C** |
| EPHB6_RAT | 481 | 484 | 4 + |  | pattern:Sx[ILV]P | . | E |
| EPHB6_RAT | 579 | 582 | 4 + |  | pattern:Sx[ILV]P | . | E |
| FLRT2_RAT | 172 | 175 | 4 + |  | pattern:Sx[ILV]P | . | E |
| FLRT2_RAT | 403 | 406 | 4 + |  | pattern:Sx[ILV]P | . | E |
| **FLRT2_RAT** | **650** | **653** | **4 +** |  | **pattern:Sx[ILV]P** | . | **C** |
| GABR1_RAT | 752 | 755 | 4 + |  | pattern:Sx[ILV]P | . | E |
| NRX2A_RAT | 875 | 878 | 4 + |  | pattern:Sx[ILV]P | . | E |
| NRX2A_RAT | 1053 | 1056 | 4 + |  | pattern:Sx[ILV]P | . | E |
| NRX2A_RAT | 1506 | 1509 | 4 + |  | pattern:Sx[ILV]P | . | E |
| **TEN2_RAT** | **178** | **181** | **4 +** |  | **pattern:Sx[ILV]P** | . | **C** |
| **TEN2_RAT** | **309** | **312** | **4 +** |  | **pattern:Sx[ILV]P** | . | **C** |
| TEN2_RAT | 1090 | 1093 | 4 + |  | pattern:Sx[ILV]P | . | E |

**Table S4.**
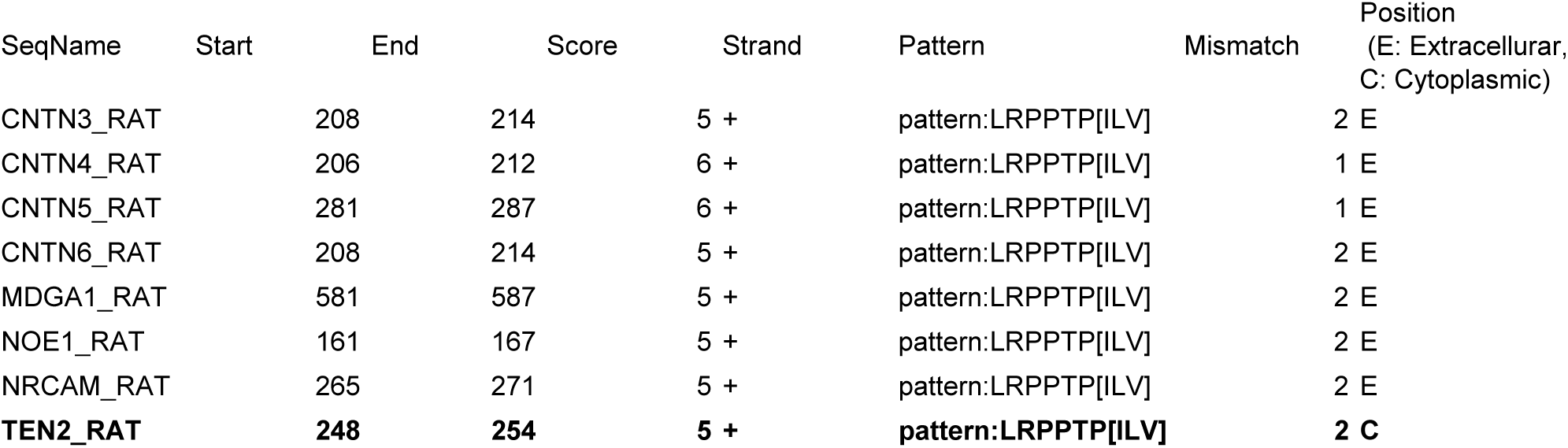
The excitatory synaptic cleft proteins with LxxPTPφ motif related to Figure 1

**Table S5.**
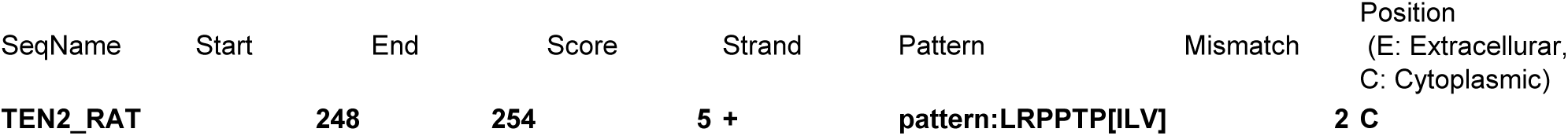
The inhibitory synaptic cleft proteins with LxxPTPφ motif related to Figure 1

**Table S6.**
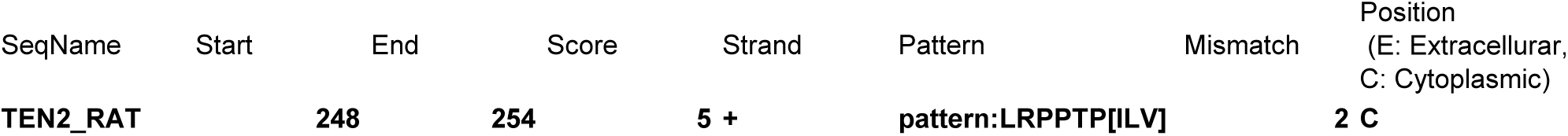
The excitatory and inhibitory common synaptic cleft proteins with LxxPTPφ motif related to Figure 1

**Table S7.** Antibody List Related to Materials and Methods

| Antibody | Source | Identifier |
| --- | --- | --- |
| anti-TEN2 Cytoplasmic | This study | N/A |
| anti-EB1 (5/EB1) | BD Biosciences | Cat# 610534, RRID:AB_397891 |
| GABRA1 antibody | Proteintech | Cat# 12410-1-AP, RRID:AB_2108692 |
| Anti-GABA-A receptor alpha5 | Synaptic Systems | Cat# 224 503, RRID:AB_2619944 |
| Anti-GABA-A receptor gamma2 | Synaptic Systems | Cat# 224 003, RRID:AB_2263066 |
| anti-gephyrin (mAb7a) | Synaptic Systems | Cat# 147011, RRID:AB_887717 |
| Anti-Bassoon | Synaptic Systems | Cat# 141 016, RRID:AB_2661779 |
| Anti-Neurologin 2 | Synaptic Systems | Cat# 129 203, RRID:AB_993014 |
| anti-PSD95 (7E3) | Cell Signaling Technology | Cat# 36233, RRID:AB_2721262 |
| anti-HA-tag (C29F4) | Cell Signaling Technology | Cat# 3724, RRID:AB_1549585 |
| Anti-IGSF9B | Merck | Cat# HPA010802, RRID:AB_1079194 |
| anti-α tubulin (DM1A) | Merck | Cat# T9026, RRID:AB_477593 |
| anti-β tubulin (TUJ1) | Covance | Cat# MMS-435P, RRID:AB_2313773 |
| anti-MAP2 | Novus | Cat# NB300-213, RRID:AB_2138178 |
| Donkey Anti-Mouse IgG (H+L), Alexa Fluor 488 | Jackson ImmunoResearch Labs | Cat# 715-546-151, RRID:AB_2340850 |
| Donkey Anti-Mouse IgG (H+L), Rhodamine Red-X | Jackson ImmunoResearch Labs | Cat# 715-296-151, RRID:AB_2340835 |
| Donkey Anti-Rabbit IgG (H+L), DyLight 405 | Jackson ImmunoResearch Labs | Cat# 711-475-152, RRID:AB_2340616 |
| Donkey Anti-Rabbit IgG (H+L), Alexa Fluor 488 | Jackson ImmunoResearch Labs | Cat# 711-546-152, RRID:AB_2340619 |
| Donkey Anti-Rabbit IgG (H+L), CF568 | Biotium | Cat# 20098-1, RRID:AB_10853318 |
| Donkey Anti-Chicken IgY (IgG) (H+L), Alexa Fluor 647 | Jackson ImmunoResearch Labs | Cat# 703-605-155, RRID:AB_2340379 |

## References

1. Aiken, J., & Holzbaur, E. L. F. (2021). Cytoskeletal regulation guides neuronal trafficking to effectively supply the synapse. Curr Biol, 31(10), R633–R650. https://doi.org/10.1016/j.cub.2021.02.024

2. Akhmanova, A., & Steinmetz, M. O. (2015). Control of microtubule organization and dynamics: two ends in the limelight. Nat Rev Mol Cell Biol, 16(12), 711–726. https://doi.org/10.1038/nrm4084

3. Beckmann, J., Schubert, R., Chiquet-Ehrismann, R., & Müller, D. J. (2013). Deciphering Teneurin Domains That Facilitate Cellular Recognition, Cell–Cell Adhesion, and Neurite Outgrowth Using Atomic Force Microscopy-Based Single-Cell Force Spectroscopy. Nano Letters, 13(6), 2937–2946. https://doi.org/10.1021/nl4013248

4. Berns, D. S., Denardo, L. A., Pederick, D. T., & Luo, L. (2018). Teneurin-3 controls topographic circuit assembly in the hippocampus. Nature, 554(7692), 328–333. https://doi.org/10.1038/nature25463

5. Charrier, C., Ehrensperger, M. V., Dahan, M., Lévi, S., & Triller, A. (2006). Cytoskeleton regulation of glycine receptor number at synapses and diffusion in the plasma membrane. J Neurosci, 26(33), 8502–8511. https://doi.org/10.1523/JNEUROSCI.1758-06.2006

6. Dahan, M., Lévi, S., Luccardini, C., Rostaing, P., Riveau, B., & Triller, A. (2003). Diffusion dynamics of glycine receptors revealed by single-quantum dot tracking. Science, 302(5644), 442–445. https://doi.org/10.1126/science.1088525

7. Del Toro, D., Carrasquero-Ordaz, M. A., Chu, A., Ruff, T., Shahin, M., Jackson, V. A., … Seiradake, E. (2020). Structural Basis of Teneurin-Latrophilin Interaction in Repulsive Guidance of Migrating Neurons. Cell, 180(2), 323–339.e319. https://doi.org/10.1016/j.cell.2019.12.014

8. Doudna, J. A., & Charpentier, E. (2014). Genome editing. The new frontier of genome engineering with CRISPR-Cas9. Science, 346(6213), 1258096. https://doi.org/10.1126/science.1258096

9. Fuhrmann, J. C., Kins, S., Rostaing, P., El Far, O., Kirsch, J., Sheng, M., … Kneussel, M. (2002). Gephyrin Interacts with Dynein Light Chains 1 and 2, Components of Motor Protein Complexes. The Journal of Neuroscience, 22(13), 5393–5402. https://doi.org/10.1523/jneurosci.22-13-05393.2002

10. Giesemann, T., Schwarz, G., Nawrotzki, R., Berhörster, K., Rothkegel, M., Schlüter, K., … Jockusch, B. M. (2003). Complex Formation between the Postsynaptic Scaffolding Protein Gephyrin, Profilin, and Mena: A Possible Link to the Microfilament System. The Journal of Neuroscience, 23(23), 8330–8339. https://doi.org/10.1523/jneurosci.23-23-08330.2003

11. Gu, J., Firestein, B. L., & Zheng, J. Q. (2008). Microtubules in dendritic spine development. J Neurosci, 28(46), 12120–12124. https://doi.org/10.1523/JNEUROSCI.2509-08.2008

12. Gu, Y., Chiu, S.-L., Liu, B., Wu, P.-H., Delannoy, M., Lin, D.-T., … Huganir, R. L. (2016). Differential vesicular sorting of AMPA and GABA_A_ receptors. Proceedings of the National Academy of Sciences, 113(7), E922–E931. https://doi.org/10.1073/pnas.1525726113

13. Guedes-Dias, P., Nirschl, J. J., Abreu, N., Tokito, M. K., Janke, C., Magiera, M. M., & Holzbaur, E. L. F. (2019). Kinesin-3 Responds to Local Microtubule Dynamics to Target Synaptic Cargo Delivery to the Presynapse. Curr Biol, 29(2), 268–282.e268. https://doi.org/10.1016/j.cub.2018.11.065

14. Gulley, R. L., & Reese, T. S. (1981). Cytoskeletal organization at the postsynaptic complex. Journal of Cell Biology, 91(1), 298–302. https://doi.org/10.1083/jcb.91.1.298

15. Heo, S., Diering, G. H., Na, C. H., Nirujogi, R. S., Bachman, J. L., Pandey, A., & Huganir, R. L. (2018). Identification of long-lived synaptic proteins by proteomic analysis of synaptosome protein turnover. Proc Natl Acad Sci U S A, 115(16), E3827–E3836. https://doi.org/10.1073/pnas.1720956115

16. Honnappa, S., Gouveia, S. M., Weisbrich, A., Damberger, F. F., Bhavesh, N. S., Jawhari, H., … Steinmetz, M. O. (2009). An EB1-binding motif acts as a microtubule tip localization signal. Cell, 138(2), 366–376. https://doi.org/10.1016/j.cell.2009.04.065

17. Hu, X., Viesselmann, C., Nam, S., Merriam, E., & Dent, E. W. (2008). Activity-dependent dynamic microtubule invasion of dendritic spines. J Neurosci, 28(49), 13094–13105. https://doi.org/10.1523/JNEUROSCI.3074-08.2008

18. Ichinose, S., Ogawa, T., & Hirokawa, N. (2015). Mechanism of Activity-Dependent Cargo Loading via the Phosphorylation of KIF3A by PKA and CaMKIIa. Neuron, 87(5), 1022–1035. https://doi.org/10.1016/j.neuron.2015.08.008

19. Ichinose, S., Ogawa, T., Jiang, X., & Hirokawa, N. (2019). The Spatiotemporal Construction of the Axon Initial Segment via KIF3/KAP3/TRIM46 Transport under MARK2 Signaling. Cell Rep, 28(9), 2413–2426.e2417. https://doi.org/10.1016/j.celrep.2019.07.093

20. Jaworski, J., Kapitein, L. C., Gouveia, S. M., Dortland, B. R., Wulf, P. S., Grigoriev, I., … Hoogenraad, C. C. (2009). Dynamic microtubules regulate dendritic spine morphology and synaptic plasticity. Neuron, 61(1), 85–100. https://doi.org/10.1016/j.neuron.2008.11.013

21. Kaneko, R., Kakinuma, T., Sato, S., & Jinno-Oue, A. (2018). Freezing sperm in short straws reduces storage space and allows transport in dry ice. J Reprod Dev, 64(6), 541–545. https://doi.org/10.1262/jrd.2018-100

22. Kaneko, T. (2017). Genome Editing in Mouse and Rat by Electroporation. Methods Mol Biol, 1630, 81–89. https://doi.org/10.1007/978-1-4939-7128-2_7

23. Kittler, J. T., Delmas, P., Jovanovic, J. N., Brown, D. A., Smart, T. G., & Moss, S. J. (2000). Constitutive Endocytosis of GABA_A_Receptors by an Association with the Adaptin AP2 Complex Modulates Inhibitory Synaptic Currents in Hippocampal Neurons. The Journal of Neuroscience, 20(21), 7972–7977. https://doi.org/10.1523/jneurosci.20-21-07972.2000

24. Kumar, A., Manatschal, C., Rai, A., Grigoriev, I., Degen, M. S., Jaussi, R., … Steinmetz, M. O. (2017). Short Linear Sequence Motif LxxPTPh Targets Diverse Proteins to Growing Microtubule Ends. Structure, 25(6), 924–932.e924. https://doi.org/10.1016/j.str.2017.04.010

25. Labonté, D., Thies, E., & Kneussel, M. (2014). The kinesin KIF21B participates in the cell surface delivery of γ2 subunit-containing GABAA receptors. Eur J Cell Biol, 93(8-9), 338–346. https://doi.org/10.1016/j.ejcb.2014.07.007

26. Li, J., Shalev-Benami, M., Sando, R., Jiang, X., Kibrom, A., Wang, J., … Araç, D. (2018). Structural Basis for Teneurin Function in Circuit-Wiring: A Toxin Motif at the Synapse. Cell, 173(3), 735–748.e715. https://doi.org/10.1016/j.cell.2018.03.036

27. Linsalata, A. E., Chen, X., Winters, C. A., & Reese, T. S. (2014). Electron tomography on γ-aminobutyric acid-ergic synapses reveals a discontinuous postsynaptic network of filaments. Journal of Comparative Neurology, 522(4), 921–936. https://doi.org/10.1002/cne.23453

28. Liu, Y.-T., Tao, C.-L., Zhang, X., Xia, W., Shi, D.-Q., Qi, L., … Bi, G.-Q. (2020). Mesophasic organization of GABAA receptors in hippocampal inhibitory synapses. Nature Neuroscience, 23(12), 1589–1596. https://doi.org/10.1038/s41593-020-00729-w

29. Loh, K. H., Stawski, P. S., Draycott, A. S., Udeshi, N. D., Lehrman, E. K., Wilton, D. K., … Ting, A. Y. (2016). Proteomic Analysis of Unbounded Cellular Compartments: Synaptic Clefts. Cell, 166(5), 1295–1307.e1221. https://doi.org/10.1016/j.cell.2016.07.041

30. Maas, C., Tagnaouti, N., Loebrich, S., Behrend, B., Lappe-Siefke, C., & Kneussel, M. (2006). Neuronal cotransport of glycine receptor and the scaffold protein gephyrin. J Cell Biol, 172(3), 441–451. https://doi.org/10.1083/jcb.200506066

31. McVicker, D. P., Awe, A. M., Richters, K. E., Wilson, R. L., Cowdrey, D. A., Hu, X., … Dent, E. W. (2016). Transport of a kinesin-cargo pair along microtubules into dendritic spines undergoing synaptic plasticity. Nat Commun, 7, 12741. https://doi.org/10.1038/ncomms12741

32. Monroy, B. Y., Tan, T. C., Oclaman, J. M., Han, J. S., Simó, S., Niwa, S., … Ori-Mckenney, K. M. (2020). A Combinatorial MAP Code Dictates Polarized Microtubule Transport. Developmental Cell, 53(1), 60–72.e64. https://doi.org/10.1016/j.devcel.2020.01.029

33. Mosca, T. J., Hong, W., Dani, V. S., Favaloro, V., & Luo, L. (2012). Trans-synaptic Teneurin signalling in neuromuscular synapse organization and target choice. Nature, 484(7393), 237–241. https://doi.org/10.1038/nature10923

34. Nakajima, K., Yin, X., Takei, Y., Seog, D. H., Homma, N., & Hirokawa, N. (2012). Molecular motor KIF5A is essential for GABA(A) receptor transport, and KIF5A deletion causes epilepsy. Neuron, 76(5), 945–961. https://doi.org/10.1016/j.neuron.2012.10.012

35. Nakata, T., Niwa, S., Okada, Y., Perez, F., & Hirokawa, N. (2011). Preferential binding of a kinesin-1 motor to GTP-tubulin-rich microtubules underlies polarized vesicle transport. J Cell Biol, 194(2), 245–255. https://doi.org/10.1083/jcb.201104034

36. O’Sullivan, M. L., de Wit, J., Savas, J. N., Comoletti, D., Otto-Hitt, S., Yates, J. R., & Ghosh, A. (2012). FLRT proteins are endogenous latrophilin ligands and regulate excitatory synapse development. Neuron, 73(5), 903–910. https://doi.org/10.1016/j.neuron.2012.01.018

37. Pawson, C., Eaton, B. A., & Davis, G. W. (2008). Formin-dependent synaptic growth: evidence that Dlar signals via Diaphanous to modulate synaptic actin and dynamic pioneer microtubules. J Neurosci, 28(44), 11111–11123. https://doi.org/10.1523/JNEUROSCI.0833-08.2008

38. Poulopoulos, A., Aramuni, G., Meyer, G., Soykan, T., Hoon, M., Papadopoulos, T., … Varoqueaux, F. (2009). Neuroligin 2 Drives Postsynaptic Assembly at Perisomatic Inhibitory Synapses through Gephyrin and Collybistin. Neuron, 63(5), 628–642. https://doi.org/10.1016/j.neuron.2009.08.023

39. Qu, X., Kumar, A., Blockus, H., Waites, C., & Bartolini, F. (2019). Activity-Dependent Nucleation of Dynamic Microtubules at Presynaptic Boutons Controls Neurotransmission. Curr Biol, 29(24), 4231–4240.e4235. https://doi.org/10.1016/j.cub.2019.10.049

40. Rice, P., Longden, I., & Bleasby, A. (2000). EMBOSS: the European Molecular Biology Open Software Suite. Trends Genet, 16(6), 276–277. https://doi.org/10.1016/s0168-9525(00)02024-2

41. Rubin, B. P., Tucker, R. P., Brown-Luedi, M., Martin, D., & Chiquet-Ehrismann, R. (2002). Teneurin 2 is expressed by the neurons of the thalamofugal visual system in situ and promotes homophilic cell-cell adhesion in vitro. Development, 129(20), 4697–4705.

42. Sando, R., Jiang, X., & Südhof, T. C. (2019). Latrophilin GPCRs direct synapse specificity by coincident binding of FLRTs and teneurins. Science, 363(6429). https://doi.org/10.1126/science.aav7969

43. Sando, R., & Südhof, T. C. (2021). Latrophilin GPCR signaling mediates synapse formation. Elife, 10. https://doi.org/10.7554/eLife.65717

44. Sanes, J. R., & Zipursky, S. L. (2020). Synaptic Specificity, Recognition Molecules, and Assembly of Neural Circuits. Cell, 181(6), 1434–1435. https://doi.org/10.1016/j.cell.2020.05.046

45. Silva, J. P., Lelianova, V. G., Ermolyuk, Y. S., Vysokov, N., Hitchen, P. G., Berninghausen, O., … Ushkaryov, Y. A. (2011). Latrophilin 1 and its endogenous ligand Lasso/teneurin-2 form a high-affinity transsynaptic receptor pair with signaling capabilities. Proc Natl Acad Sci U S A, 108(29), 12113–12118. https://doi.org/10.1073/pnas.1019434108

46. Skube, S. B., Chaverri, J. M., & Goodson, H. V. (2010). Effect of GFP tags on the localization of EB1 and EB1 fragments in vivo. Cytoskeleton (Hoboken), 67(1), 1–12. https://doi.org/10.1002/cm.20409

47. Söderberg, O., Gullberg, M., Jarvius, M., Ridderstråle, K., Leuchowius, K. J., Jarvius, J., … Landegren, U. (2006). Direct observation of individual endogenous protein complexes in situ by proximity ligation. Nat Methods, 3(12), 995–1000. https://doi.org/10.1038/nmeth947

48. Takács, V. T., Freund, T. F., & Nyiri, G. (2013). Neuroligin 2 Is Expressed in Synapses Established by Cholinergic Cells in the Mouse Brain. PLoS ONE, 8(9), e72450. https://doi.org/10.1371/journal.pone.0072450

49. Twelvetrees, A. E., Lesept, F., Holzbaur, E. L. F., & Kittler, J. T. (2019). The adaptor proteins HAP1a and GRIP1 collaborate to activate kinesin-1 isoform KIF5C. Journal of Cell Science, 132(24), jcs215822. https://doi.org/10.1242/jcs.215822

50. Twelvetrees, A. E., Yuen, E. Y., Arancibia-Carcamo, I. L., MacAskill, A. F., Rostaing, P., Lumb, M. J., … Kittler, J. T. (2010). Delivery of GABAARs to synapses is mediated by HAP1-KIF5 and disrupted by mutant huntingtin. Neuron, 65(1), 53–65. https://doi.org/10.1016/j.neuron.2009.12.007

51. Uchigashima, M., Ohtsuka, T., Kobayashi, K., & Watanabe, M. (2016). Dopamine synapse is a neuroligin-2– mediated contact between dopaminergic presynaptic and GABAergic postsynaptic structures. Proceedings of the National Academy of Sciences, 113(15), 4206–4211. https://doi.org/10.1073/pnas.1514074113

52. Vysokov, N. V., Silva, J. P., Lelianova, V. G., Suckling, J., Cassidy, J., Blackburn, J. K., … Ushkaryov, Y. A. (2018). Proteolytically released Lasso/teneurin-2 induces axonal attraction by interacting with latrophilin-1 on axonal growth cones. Elife, 7. https://doi.org/10.7554/eLife.37935

53. Woo, J., Kwon, S.-K., Nam, J., Choi, S., Takahashi, H., Krueger, D., … Kim, E. (2013). The adhesion protein IgSF9b is coupled to neuroligin 2 via S-SCAM to promote inhibitory synapse development. Journal of Cell Biology, 201(6), 929–944. https://doi.org/10.1083/jcb.201209132

